# Optimizing spectral quality with quantum dots to enhance crop yield in controlled environments

**DOI:** 10.1101/2020.06.17.157487

**Authors:** Charles H. Parrish, Damon Hebert, Aaron Jackson, Karthik Ramasamy, Hunter McDaniel, Gene A. Giacomelli, Matthew R. Bergren

**Affiliations:** Controlled Environment Agriculture Center, University of Arizona, Tucson, Arizona 85719, USA; UbiQD, Inc., Los Alamos, New Mexico 87544, USA

## Abstract

Bioregenerative life-support systems (BLSS) involving plants will be required to realize self-sustaining human settlements beyond Earth. To improve plant photosynthetic efficiency in BLSS, the quality of the solar spectrum can be modified by lightweight, luminescent films. CuInS_2_/ZnS quantum dot (QD) films with peak emissions at 600 and 660 nm were used to redshift ultraviolet and blue photons to increase biomass accumulation in red romaine lettuce. Plant growth parameters, except spectral quality, were held constant among three equivalent production environments. Lettuce grown under the 600 and 660 nm-emitting QD films respectively increased edible dry mass (13% and 9%), edible fresh mass (11% each), and total leaf area (8% and 13%) compared to under a control film that contained no QDs. Spectral modifications by the luminescent QD films improved photosynthetic efficiency in lettuce and could enhance crop productivity in greenhouses on Earth or in space, where further improvements are expected from greater availability of ultraviolet photons for conversion.

## Introduction

As anthropogenic climate change exacerbates the severity and frequency of extreme weather events,^1^ arable farmland is decreasing due to overpopulation and drought.^2^ For resilience against these threats, producers of vegetable crops have turned to controlled environment agriculture (CEA) as a solution. CEA affords crop production with the highest resource use efficiency (RUE), and exceeds the annual per-area yield of field agriculture ten-fold.^3^ This greater productivity results from optimized environmental controls, year-round production, and reduced pests and diseases. Furthermore, recirculating hydroponic plant nutrient delivery systems recycle water for greatest RUE.^4^

CEA food production is particularly advantageous in extreme environments, like outer space, where regular access to fresh vegetables is impractical. NASA and other space agencies are developing CEA systems for crewed space applications, which range from growth chambers with light-emitting diode (LED)-based lighting to drive photosynthesis,^5^ to deployable greenhouses that could utilize sunlight for photosynthesis, like the Prototype Mars-Lunar Greenhouse (MLGH).^6^

Until recently, no viable options were available for controlling the quality of light for greenhouse crop production except for expensive and energy-intensive electrical lighting.^7^ Furthermore, artificial lighting requires relatively heavy electrical equipment to deploy, operate, and maintain, which limits applications for extended space missions. Therefore, technologies that improve light quality without increasing energy demand, excessive physical mass, deployment challenges, or expense, would be beneficial for terrestrial horticulture and are required for space applications where these constraints are even greater.

A new technology that uses luminescent CuInS_2_/ZnS (CIS/ZnS) quantum dots (QD) embedded in flexible films to passively modify the solar spectrum is presented which can improve light quality and thereby enhances the photosynthetic efficiency of plants grown in CEA.

## Results

### Spectral Quality on Plant Growth

The 400 to 700 nm waveband of the electromagnetic spectrum primarily powering photosynthesis is referred to as photosynthetically active radiation (PAR). The amount of PAR photons a plant receives is quantified by the photosynthetic photon flux density (PPFD, μmol m^-2^ s^-1^), which is the number of photosynthetically active photons per unit area per second. The PPFD a plant receives in a 24-hour day is directly proportional to the rate of plant growth, determined as biomass accumulation which is measured by the daily light integral (DLI, mol m^-2^ d^-1^). While PPFD and DLI are important plant growth parameters for characterizing plant development, they ignore spectral quality differences of radiation received by the plant.

Spectral quality is an important growth parameter since the efficiency of photosynthesis is dependent on the wavelength of the photon absorbed by the plant. The wavelength-dependent photosynthetic efficiency can be determined by the photosynthetic quantum yield (QY), a measure of the production of oxygen or consumption of carbon dioxide, for various crops.^8,9^ The photosynthetic action spectrum (Figure 1a, lettuce), compares the relative photosynthetic QY per wavelength, and indicates that wavelengths between 575 nm and 675 nm are ∼30% more efficient for photosynthesis than photons in the blue waveband (400-500nm).

**Figure 1.**
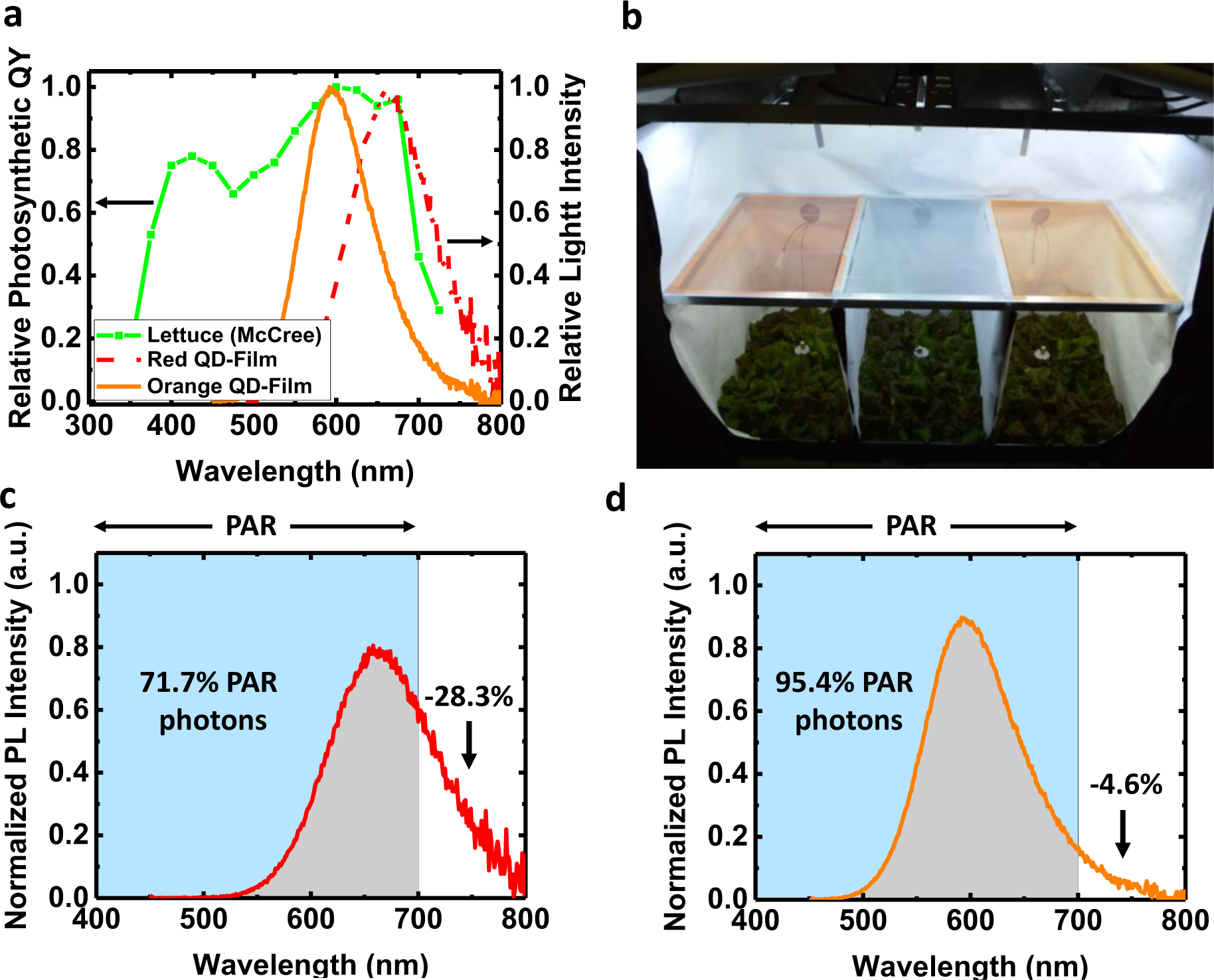
Luminescent properties of QD films. (a) The emission spectra of the two QD films with peak PL emissions of 600 nm (orange, solid line) and 660 nm (red, dashed line) overlapped on the photosynthetic response curve for lettuce (green, squares) measured by McCree.^8^ (b) Photograph of the PGTC for lettuce plant trials. Three light treatment areas were used to compare different light spectra created by QD films (left and right) to a Control film (middle) under similar light intensity. The spectra of QD films with peak emission of (c) 660 nm and (d) 600 nm were normalized to have an area of 1 and then integrated over PAR wavelengths (400-700 nm) to determine the portion of the emission that was outside of PAR.

Several recent studies about light spectra effects on plant growth have determined that both light quantity and quality affected plant morphology.^10-13^ Using a broad-spectrum light source to grow plants outperformed monochromatic light treatments, but modifying the relative percentages of different wavelengths did produce varied and improved results. For example, the sensitivity of seven plant species to blue and green light with a daylight-equivalent 500 µmol m^-2^ s^-1^ PPFD, reduced dry mass in tomatoes cucumbers, and peppers when blue light was increased between 11% and 28% of the total spectrum^.^13^^ At 200 µmol m^-2^ s^-1^ PPFD, tomato continued to be negatively affected, where dry mass was reduced by 41%, while no difference was observed in pepper or cucumber. These results illustrated the importance of spectral quality for influencing plant response, particularly in high light intensity environments, and suggest that there are optimal spectra for different plant species.

### Spectral control with CIS/ZnS QD films

QDs have been utilized for a variety of applied technologies, including remote phosphors for displays,^14,15^ solid-state lighting,^16,17^ solar harvesting electricity generation,^18,19^ cancer detection,^20-22^ and even for studying pollination habits of bees.^23,24^ Here we introduce a new application, incorporating QDs into luminescent agriculture films to improve crop production in greenhouses. Luminescent QD films enable passive modification of the solar spectrum to improve light quality. In this study, CIS/ZnS QDs are incorporated into a film to down-convert UV and blue photons to orange and red photons. Compared with other QD compositions, CIS/ZnS QDs have unique optical properties: a size-tunable PL emission, a very high PL quantum yield (QY), and a broad spectrum emission.^25^ For QDs in general, a broad emission is generally associated with a dispersion of QD sizes or compositions in an ensemble and typically result in lower conversion efficiencies due to an unoptimized synthesis procedure, but for CIS/ZnS this is a result of a defect-mediated emission.^26^ Additionally, CIS/ZnS QDs exhibit a large Stokes shift, allowing for absorption in the UV and blue (see Supplemental Information) while emitting in the red or orange, which minimizes PAR absorption and shift the spectrum to more photosynthetically efficient wavelengths.

Two different QD films were manufactured, with peak emissions centered at 600 nm (Orange, O-QD) and 660 nm (Red, R-QD). These wavelengths were chosen due to the strong overlap between their emission spectra and the wavelength range associated with the highest relative photosynthetic QY for lettuce (Figure 1a).

To evaluate how the modified spectra from the QD films affect plant growth, indoor plant trials were conducted with red romaine lettuce (*Lactuca sativa* L. cv. ‘Outredgeous’) in a custom-built, plant growth test chamber (PGTC). The PGTC was designed to maintain a uniform environment across all plant growing areas, two of which were under O-QD and R-QD films and the third was under a Control (C) polyethylene film, without QDs (Figure 1b).

Commercially available horticultural luminaires and metal-halide (MH) lamps were installed above the three horizontally suspended films (see Methods). The light passing through the films to the plant canopies below provided spatially uniform illumination of approximately 380 μmol m^-2^ s^-1^ PPFD and had a spectrum that was similar to the solar spectrum (see Supplemental Information). Other plant growth parameters, such as the air and root temperatures, atmospheric CO_2_ concentration, PPFD, pH, and electrical conductivity (EC) were monitored and remained equivalent for all three treatment areas in each plant trial (see Supplemental Information). This ensured that only the quality of the light spectrum was varied between each treatment area.

The spectra under the O-QD and R-QD films were measured with a spectroradiometer (Apogee Instruments, PS-300) and compared with the spectra under the C film. The total PPFD and the spectral distribution within each light treatment zone is shown in Table 1, The ultraviolet (UV) and blue (B) wavelengths were clearly reduced under both of the QD films compared to the control, while the intensities of O, R, and far-red (FR) were increased, compared with the control.

**Table 1.** Spectral distribution under QD and control films and light treatment provided to the plants.

| **Film** | **PPFD**<br>($\mu\text{mol m}^{-2} \text{s}^{-1}$) | **UV**<br>($<400 \text{ nm}$) | **B**<br>($400\text{-}500 \text{ nm}$) | **G**<br>($500\text{-}600 \text{ nm}$) | **R**<br>($600\text{-}700 \text{ nm}$) | **FR**<br>($700\text{-}800 \text{ nm}$) |
| --- | --- | --- | --- | --- | --- | --- |
| Control | 382 | 2.1% | 31.3% | 35.8% | 20.6% | 10.2% |
| Orange | 395 | 1.4% | 23.3% | 36.4% | 26.7% | 12.1% |
| Red | 338 | 0.9% | 20.5% | 32.3% | 29.1% | 17.2% |

To ensure differences in production were solely resulting from spectral differences, the light intensity for each treatment area were equal. While the difference between PPFD values measured under the C and O-QD films were within 3%, PPFD under the R-QD film the PPFD was 12% lower than the control film. This lower value was attributed to a portion of the R-QD emission extending beyond 700 nm, which is outside the PAR range, and therefore was not measured by the PAR sensor. This is illustrated in Figure 1c and 1d, where normalized emission spectra of the R-QD and O-QD films, respectively indicated only 4.6% of the emission spectrum from O-QD film was outside of PAR (>700 nm), but 28.3% from the R-QD film did not contribute to PPFD. Therefore, even if the total number of incident photons was held constant for the three treatment areas, the measured PPFD would always be lower for the R-QD film compared to the other light treatments. The concentration of QDs in the O-QD and the R-QD films were similar and designed to ensure similar overall absorption of the MH light source. Since both films had similar absorption and had equivalent PL QYs of ∼85%, equivalent quantities of photons were converted by each film. With a spectroradiometer the spectral quality from 300 to 850 nm was also measured to evaluate changes in film performance throughout the experiment. No spectral differences were observed over the three experiments (Figure 2a and 2b).

**Figure 2.**
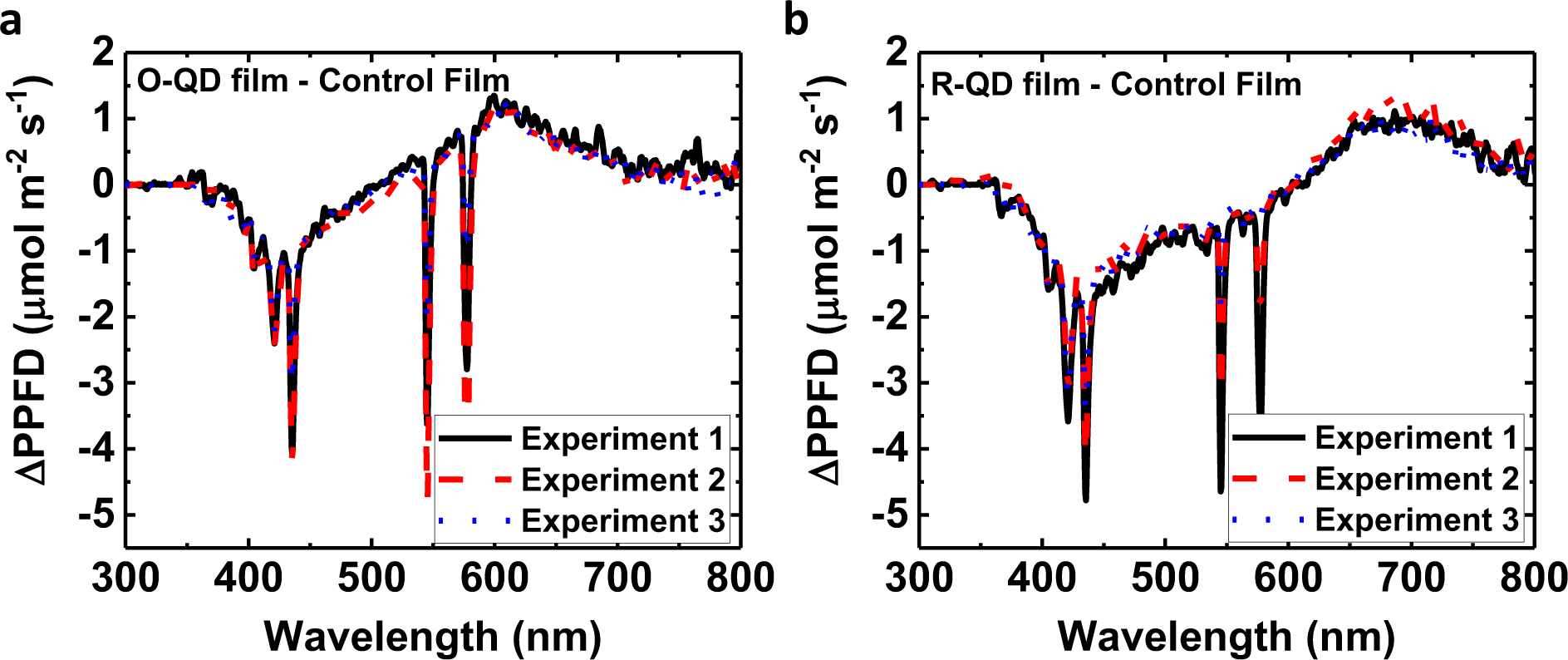
Changes in spectral differences of QD-films over time. Spectral differences (ΔPPFD) between each QD-film and control film were recorded before each experiment. Using a spectroradiometer the spectrum measured under the O QD-film (a) and R QD-film (b) were subtracted from the spectrum measured under the C film before each experiment.

### Spectral effects on red romaine lettuce production

The plant study consisted of three, 28-day experiments consisting of 36 lettuce plants (N = 36) divided equally among three experimental treatments (n = 12). Plants were harvested 28 days after sowing (DAS). Post-harvest measurements included total leaf area (TLA, cm^2^), edible fresh mass (FM, g), and edible dry mass (DM, g) and are shown in Figure 3). TLA was calculated with image recognition software, and manual corrections were performed as needed (see Supplemental Information). The edible portions of the plant include the leaf and shoot mass, but not the root mass. A Student’s t-test ANOVA was performed on the model that included two factors, each at three levels which tested all pairwise comparisons of the effect least squares mean. The levels included O-QD, R-QD, and Control (C) film treatments, while the second factor included replicate experiment numbers 1, 2, and 3. In cases where the constant variance assumption of the Student’s t-test was violated, logarithmic transformation of the data was conducted to analyze the data. Cohen’s d effect sizes were calculated to determine the percentage differences in edible DM, edible FM, and TLA between each QD film and C. The effect size gauged the contribution of only one factor among multiple factors (film treatment and experiment number), and as such showed the significance of the contribution from the film treatment only, despite noise in the variation from combining both factors. The results from the three film treatments across all three repeated experiments are summarized in Table 2.

**Table 2.**
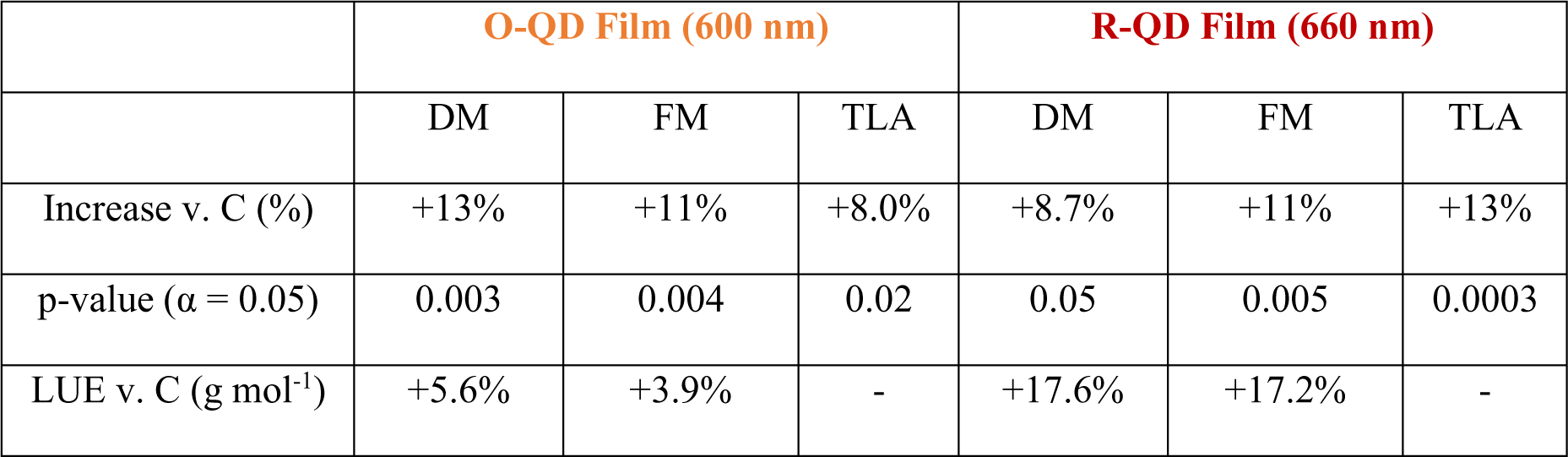
Percentage differences and significance of effect sizes, and calculated lighting-use efficiency (LUE) of O-QD and R-QD Films compared with Control (C). Statistically significant increases were observed in dry mass (DM), fresh mass (FM), and total leaf area (TLA) at confidence level α = 0.05.

|  | **O-QD Film (600 nm)** |  |  | **R-QD Film (660 nm)** |  |  |
| --- | --- | --- | --- | --- | --- | --- |
|  | DM | FM | TLA | DM | FM | TLA |
| Increase v. C (%) | +13% | +11% | +8.0% | +8.7% | +11% | +13% |
| p-value ($\alpha = 0.05$) | 0.003 | 0.004 | 0.02 | 0.05 | 0.005 | 0.0003 |
| LUE v. C ($\text{g mol}^{-1}$) | +5.6% | +3.9% | - | +17.6% | +17.2% | - |

**Figure 3.**
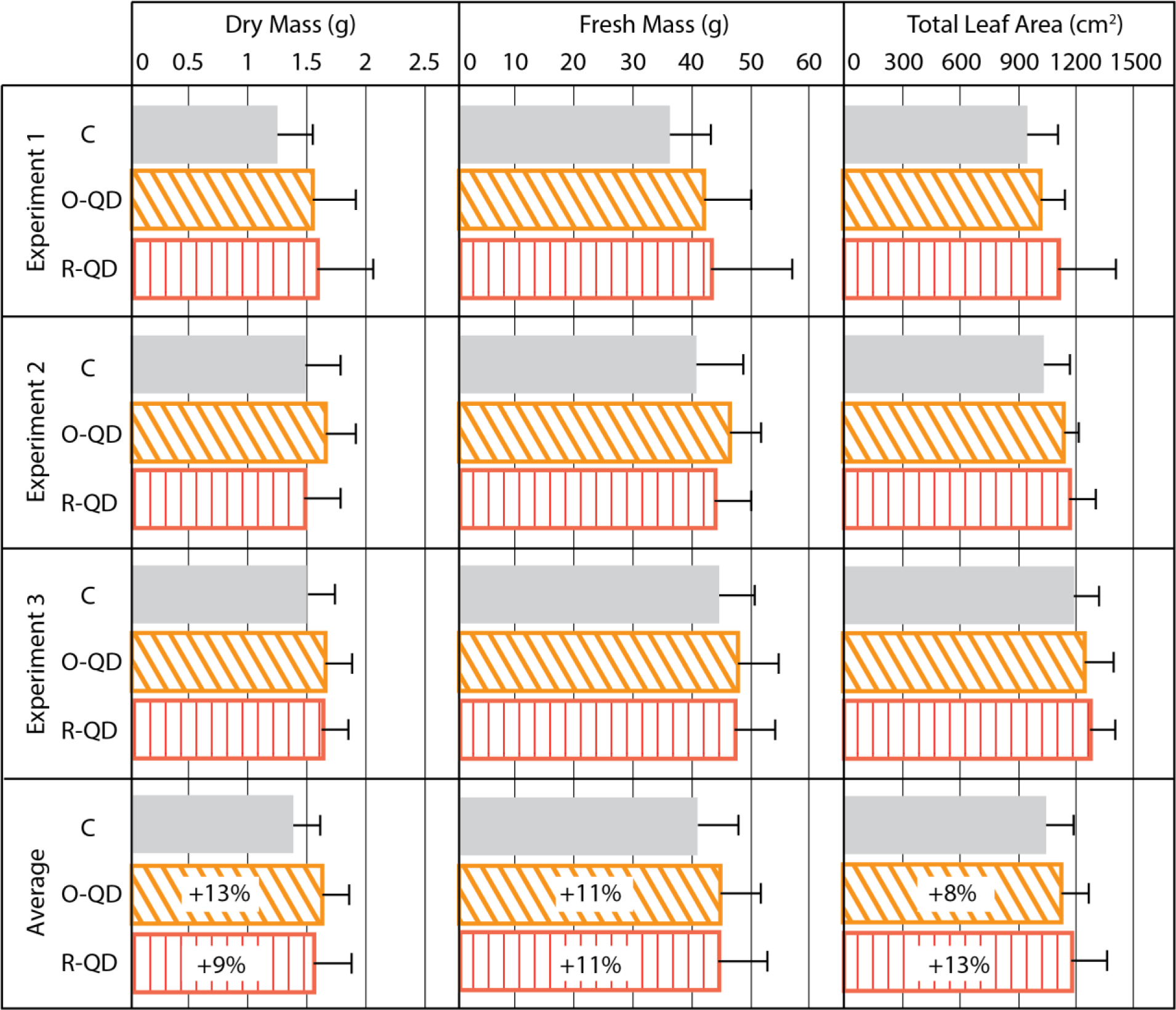
Experimental results of lettuce plant trials. Experimental results of measured edible dry mass, edible fresh mass, and total leaf area for three replicate lettuce experiments under three different light spectra (n = 12 per group in each treatment; N = 36 per experiment. Error bars show one standard deviation. Effect sizes for each metric were statistically significant with a confidence level of 95%.

These results provided statistically significant data that the QD films improved overall crop yield. The photosynthetic efficiency of the plants was improved because of light quality from the QD films, as more biomass was produced for DLI values compared to the control film. In general, when DLI is reduced, the biomass accumulation is reduced in direct proportion a 1% reduction in DLI correlates with a 1% reduction in biomass production for most plants.^27^

In these plant experiments, the opposite was observed for the R-QD treatment. Even with a reduction in DLI of 12% under the R-QD film, statistically significant increases were observed in edible DM, FM, and TLA. Thus, the light use efficiency (LUE, g mol^-1^) was greater under the R-QD film, which directly correlates with the photosynthetic efficiency of the plant. The LUE was calculated by dividing the grams of fresh or dry mass produced per day by the DLI (see Supplemental Information). Table 2 illustrates how the R-QD film and O-QD films exhibited an improved LUE compared with the control.

### QD Films for Space Applications

From the plant experiments above, it was demonstrated that modifying the solar spectrum with luminescent QD films would lead to increased plant production by improving the photosynthetic efficiency of the plant. If this technology were deployed in greenhouses on the moon or Mars, additional benefits to plant production could be realized due to the larger availability of UV photons for conversion into PAR. To estimate these benefits, a simple model described below was employed. First, the absorption spectrum of the O-QD and R-QD films (see Supplemental Information) was used to calculate the percentage of absorbed photon flux for both terrestrial and extraterrestrial locations. This was performed by convoluting each QD film absorption spectra with the solar spectrum for reference air mass (AM) 1.5 on Earth and AM 1.0 in space.^28^ With the convoluted spectra, the absorption percentage of UV (<400 nm), B (400 - 500 nm), and G (500 - 600 nm) were calculated for each film by integrating the area under the convoluted absorption curve for each wavelength range (see Figure 4).

**Figure 4.**
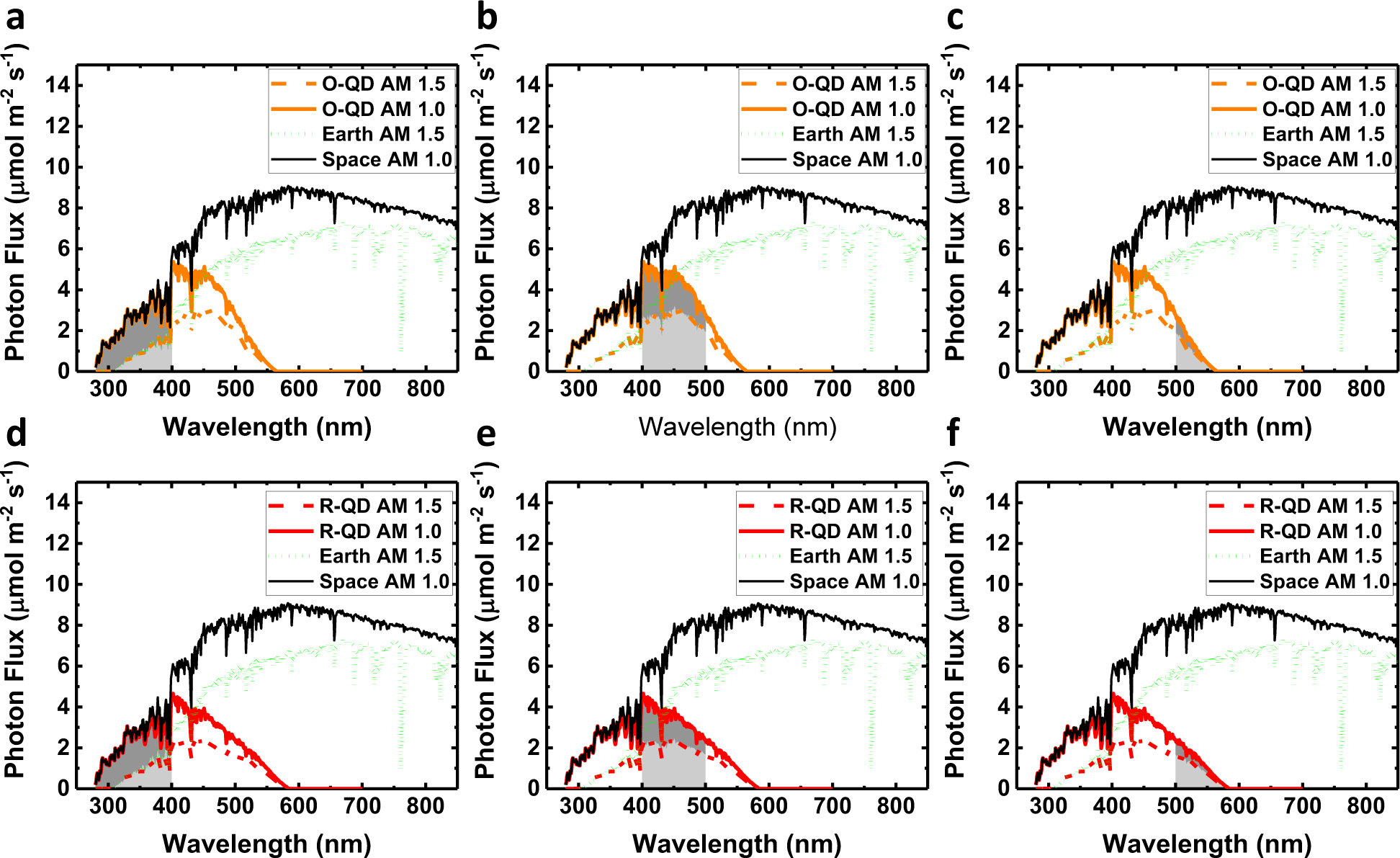
Comparison between absorbed photon flux for QD-films under AM 1.0 and AM 1.5 spectra. Convoluted absorption spectra of the O-QD film with the AM 1.0 and AM 1.5 spectra are presented in a-c, where the integrated areas of absorbed photon flux of the (a) UV (<400nm), (b) blue (400-500 nm), and (c) green (500-600 nm) spectral ranges are calculated for each solar spectra and compared. Similarly, convoluted absorption spectra of the R-QD film with the AM 1.0 and AM 1.5 spectra are presented in d-f, where the integrated areas of absorbed photon flux for the (d) UV, (e) blue, and (f) green spectral ranges are calculated for each solar spectra and compared.

After calculating how much light is absorbed for a given wavelength range, the total photon flux density converted into PAR from the emission of the QDs was calculated. As a first approximation, the following assumptions were included: 1) the QY of the films was 100%, 2) the emission spectrum of the QDs was fully within the PAR region (400 to 700 nm), and 3) all emitted photons were directed toward the plants. Considering these assumptions, the net change in PAR would simply be the addition of the absorbed UV photon flux density that is converted into PL, since the converted blue and green photons to PL would remain within PAR under these assumptions. However, loss mechanisms present in a real system are unaccounted under the above assumptions and therefore must be considered to obtain a more realistic estimate. The first loss mechanism to consider is the QY of the QD films not being 100%. In this case, both QD films had a measured PL QY of 85%. Second, due to the isotropic PL emission, it can be assumed that ∼25% of the PL would not be able to be absorbed by the plants due to PL emission in the opposite direction. Third, as mentioned previously, there are portions of the emission spectrum for both QD films that extended beyond the PAR region and thus wouldn’t contribute to PAR. Including these three loss mechanisms, the estimated net difference in PPFD contributions were calculated for the O-QD and R-QD films under both the AM 1.0 and AM 1.5 solar spectra. Table 3 summarizes the results.

**Table 3.** Estimated changes in PPFD due to absorption and emission of QD films for AM 1.0 and AM 1.5 solar spectrum

| **AM 1.0** |  |  |  |  |  |  |
| --- | --- | --- | --- | --- | --- | --- |
| **Film** | **Total PPFD**<br>**($\mu\text{mol m}^{-2} \text{s}^{-1}$)** | **$\Delta\text{PPFD}$**<br>**UV (%)** | **$\Delta\text{PPFD}$**<br>**Blue (%)** | **$\Delta\text{PPFD}$**<br>**Green (%)** | **Net $\Delta\text{PPFD}$**<br>**(%)** | **Net $\Delta\text{PPFD}$**<br>**($\mu\text{mol m}^{-2} \text{s}^{-1}$)** |
| O-QD | 2413.2 | +8.6% | -5.1% | -0.9% | +2.6% | +62.6 |
| R-QD | 2413.2 | +6.2% | -6.6% | -2.0% | -2.4% | -58.3 |
| **AM 1.5** |  |  |  |  |  |  |
| O-QD | 1735.3 | +3.7% | -4.3% | -0.9% | -1.6% | -26.8 |
| R-QD | 1735.3 | +2.6% | -5.6% | -2.0% | -5.0% | -86.0 |

For the AM 1.0 spectrum, the PPFD below the O-QD film would in theory be increased by approximately 63 µmol m^-2^ s^-1^ of photon flux density (+2.6%), while the R-QD film would have an overall decrease in PPFD (−2.4%). For the AM 1.5 spectrum, neither of the films would be expected to improve PPFD, with respective decreases of -1.6% and -5% under the O-QD and R-QD films. Comparing the net ΔPPFD for the QD films under AM 1.0 and AM 1.5 solar spectra shows that there would be a relative improvement of +4.2% in PPFD for the O-QD film and +2.6% relative improvement in PPFD for the R-QD film under the AM 1.0 spectrum.

From the calculations above, it can be concluded that for both solar spectra the net PPFD would be expected to decrease under the R-QD film, but a potential net increase in PPFD under the O-QD film would be expected for the AM 1.0 spectra. For the AM 1.5 spectra, a slight reduction in PPFD would still be expected for the O-QD film due to inherent losses in the system and to less UV light availability for shifting into PAR. In general, 1% of additional DLI would result in a 1% increase in production for most plants,^27^ so the estimated additional benefit for crop production in space applications would be 4.2% and 2.6% for the O-QD and R-QD films, respectively, due to converting more UV photons. This would be in addition to the observed increased production due to spectral quality shown in the plant experiments above. For terrestrial applications, a net increase in PPFD would not be expected under either QD film; therefore, improved production would only be expected from spectral quality.

It is noteworthy that the AM 1.0 spectrum used in these calculations was taken from the ASTM standard ASTME-490, which is based on irradiance measurements made by satellites and other equipment, and relates to the intensity at approximately the distance of Earth to the Sun.^28^ In space mission, such as to the Moon or Mars, the intensity would be reduced by ∼1/d^2^, where d is the distance from Earth to the new location and could be further reduced by a different atmosphere. Since the Moon has no atmosphere and it orbits the Earth, there wouldn’t be an expected reduction in available PPFD. However, in a location like Mars, the intensity would be reduced to ∼1/3 of the AM 1.0 spectrum measured at Earth, due to Mars being ∼1.5x further from the Sun and having an atmosphere, which further reduces the UV portion by an additional ∼4%.^29^ Since ∼39% more PPFD is available under an AM 1.0 spectrum (measured at Earth) compared to the AM 1.5 spectrum this would partially make up for the PAR reduction on Mars, but there would still be an expected overall reduction in PPFD compared to terrestrial applications. These considerations further demonstrate the importance of maximizing the LUE for the plants, as the intensity of sunlight would continue to drop as crewed space missions expand farther out into the solar system.

## Discussion

In summary, this work introduced and demonstrated the benefits of a novel luminescent film technology enabled by CIS/ZnS QDs to improve spectral quality for plant growth. The edible DM under the O-QD and R-QD films was increased by +13% and +9%, respectively, and the edible FM was increased by +10% under both films. The TLA was also improved under the O-QD and R-QD films by +8% and +13%, respectively. These results clearly indicated that lettuce grown under the QD films exhibited more efficient growth than the Control even with a DLI reduction of 12% under the R-QD film. This is supported by LUE calculations, where the DM was 18% greater under the R-QD film and 6% greater under the O-QD film. Finally, a mathematical model was implemented to estimate crop yield improvements under QD films when deployed in the AM 1.0 solar spectrum in space. The plant production improvements from these QD films would potentially increase crop yields in CEA cultivation on Earth, and, due to their relative light weight and small form factor, would be beneficial for long-duration crewed space missions.

## Methods

### Plant growth test chamber design and operation

Lettuce plant responses to the modified spectrum of light provided by the QD films were determined within a custom-built plant growth test chamber (PGTC). The PGTC framework (1.8 x 1.0 x 2.2 m, L x W x H) consisted of three platforms (from top to bottom): a lamp mounting platform located 61 cm above the test films support platform, which was 91 cm above the plant growth platform.

A nutrient storage tank was located below the plant growth platform at the base of the PGTC. The lamp platform had four P.L. Light Systems luminaires fitted with metal-halide (MH) lamps (Solistek, 400-W 10k Finisher) to provide uniform radiation distribution above the test film support frame (1.8 m x 1 m), that held three horizontally mounted and adjacent test films. The plant growth platform was comprised of a 1.8 m x 1 m root zone nutrient solution tray, shared by plants grown within each of three separate zones (60 x 86 x 91 cm, L x W x H). The zones were delineated with a white-on-black plastic film (California Grow Films ORCA Grow Film, 98% reflection and 99% diffusion of the visible spectrum) that was installed to maximize light intensity, ensure light uniformity, and separate each light treatment by eliminating cross-contamination of light between zones.

The air and root zone temperatures, atmospheric CO_2_ concentration, and nutrient solution pH and EC were continuously monitored and remained equivalent for the plants within each of the three zones, thereby having only the light quality based on the light spectrum distribution as modified by the test film as a variable (see Supplemental Information for characteristics of environmental uniformity). A ventilation fan was installed in each test zone to exchange cool, dry air from the room with the warm, moist air of the test zone and to maintain ambient CO_2_ concentration, while providing air movement among the plant leaves.

The plants were secured and spaced 12 cm x 12 cm (55 plants m^-2^, see Supplemental Information for plant layout) within a rigid, high-density polyethylene (HDPE) board (61 x 100 x 2.5 cm, L x W x H) covered with ORCA Grow Film and set on top of a water-tight basin. A modified Hoagland hydroponic plant nutrient solution (see Supplemental Information for formulation) was continually distributed and recirculated to the common root zone environment of all of the plants. The nutrient solution was maintained at a constant depth of 2.5 cm in the root zone tray by sub-irrigation from the nutrient solution storage tank (83 L), providing one volume change of solution every five minutes.

A data acquisition and control system (Campbell Scientific 21X Micrologger® datalogger) monitored and recorded 15-minute averages of plant microclimate conditions of the PGTC measured at 5-second intervals. The datalogger also activated the lamps to control the photoperiod (12:00-02:00, 14 h), and it controlled the nutrient solution pH (6.1-6.2; Weiss Research PHS-0201-3B). Quantum sensors (400-700 nm; Apogee Instruments SQ-500-SS; error ± 0.8%) and radiation-shielded thermistors (Campbell Scientific 107 BetaTherm 100K6A1IA, error ± 0.5 °C) were installed in each test zone of the PGTC to measure PPFD (μmol m^-2^ s^-1^) and air temperature (°C), respectively. Average photoperiod/scotoperiod air temperatures were 24/20 °C, respectively, and average air temperature differences between each test zone remained within ± 0.5 °C throughout the experiments. The atmospheric CO_2_ concentration (ppm; Vaisala GMT222, error ± 1.5%) was monitored in the room adjacent to the PGTC but not controlled and ranged from 361-376 ppm for each experiment.

The recirculating, deep-water culture hydroponic nutrient delivery system consisted of a pump, four emitters, one drain, and two cylindrical storage reservoirs connected with each other at both ends. The shared nutrient solution temperature (°C, Type T copper-constantan thermocouple, error ± 1.0 °C) and electrical conductivity (EC, mS cm^-1^; Hanna Instruments HI-3001) were measured and recorded. The acidity of the nutrient solution was controlled to maintain a pH setpoint (6.1-6.2) by the addition of dilute nitric acid into the nutrient storage reservoir by a peristaltic pump. The nutrient storage tank was monitored and manually refreshed daily to maintain EC setpoint (1.8-1.9 mS cm^-1^) by addition of pre-mixed nutrient solution. Nutrient solution and/or tap H2O was manually added to the storage reservoirs on a daily basis to ensure the EC remained on target. The nutrient solution was pumped up from the reservoirs to the back wall of the PGTC, where the manifold of four equidistant emitters constantly output nutrient solution to the lettuce plants in the shared basin. The nutrient solution drained from a single drain at the opposite end of the basin, where it returned to the nutrient storage reservoirs for recirculation.

### Plant studies in the PGTC

In each plant experiment, lettuce plants were established by double-seeding 150, 3.8-cm rockwool starter cubes (Grodan AO 36/40) with red romaine lettuce (*Lactuca sativa* L. cv. ‘Outredgeous’; Johnny’s Seeds, organic variety). They were placed in the three plant zones in the PGTC for 48-72 h without light till seed germination was observed. Germination was marked by root radicle emergence from the seed, after which a 14-h photoperiod (14 h of light per 24-hour period) was initiated to provide a target average daily light integral (DLI) of 17 mol m^-2^ d^-1^ (see Supplemental Information for DLI values). At 7 DAS, the first true leaves emerged, and thinning was performed to leave one healthy seedling in each rockwool cube. Extra rockwool cubes were removed from the floating raft within each of the three treatment zones to achieve a final planting density of 55 plants m^-2^. The experimental treatments continued to harvest at 28 DAS.

The rockwool cubes were irrigated continuously with pH 6.2 tap water until germination, and between germination and thinning they were irrigated with half-strength modified Hoagland’s nutrient solution. After thinning, full-strength solution was used in the recirculating hydroponic basin which continuously pumped stored nutrient solution to plants in all three zones for the duration of the experiment at a system turnover rate of 5 min.

Harvest occurred at 28 DAS when the edible fresh mass (g) was measured for each data plant (n=12 per treatment, N=36 total). Leaves were then flattened and photographed for measurement of total leaf area (TLA, cm^2^) with image analysis software (see Supplemental Information for software description). After photographing, each plant sample was individually bagged and labeled, dried in a drying oven at 40 °C for 72 h to ensure thoroughly uniform drying (see Supplemental Information for percentage dry mass values), and weighed to determine dry mass (g).

### QD film manufacture and optical properties

The CuInS^2^ (CIS)/ZnS QDs used in this study were manufactured using methods similar to those described in reference 30 with slight modifications to optimize peak wavelength emission and QY. Size of the orange (590 nm PL emission) and red (630 nm PL emission) QDs are 4.8 ± 0.7 nm and 5.1 ± 0.9 nm, respectively, and have typical size distribution of ∼15-19%. The CIS QDs were incorporated into agriculture films 60 cm x 91 cm in size by dispersing the nanoparticles into an acrylic resin (described in reference 31), which was then coated between two sheets of polyethylene terephthalate (PET) barrier film (manufactured by I-Components, Co.) using a drawdown method. The water vapor transmission rate for the PET barrier films was 0.09 (g/m^2^/day). The laminate film was then exposed to 405-nm emitting LEDs to cure the nanocomposite interlayer. The total thickness of the films was 350 µm with an interlayer thickness of 150 µm. After curing the QD resin between the two PET layers, the optical properties of both films were characterized. The PL QY was measured with a commercial PL spectrometer (Horiba Fluoromax 4) equipped with an integrating sphere and using an excitation wavelength of 440 nm. The QY of each film was measured to be 85% ± 5%, with a peak emission for the O-QD film at 600 nm (120 nm FWHM) and at 660 nm (120 nm FWHM) for the R-QD film. The haze of both QD films and the bi-layer, 6 mil polyethylene (PE) C film was measured using a custom-built setup consisting of a collimated 640-nm LED light source, a 30.5-cm diameter integrating sphere, and a fiber optic spectrometer (Avantes AvaSpec Dual Channel). The haze for all three films was measured to be 2% ± 0.5%.

## Acknowledgements

This work was funded under a Small Business Technology Transfer contract by the National Aeronautics and Space Administration under Award No. 80NSSC18P2144 and the University of Arizona ALVSCE Bridge Funding Program 2019. The total leaf area analysis program was developed and implemented by KC Shasteen, and consultation on statistical methods was provided by Lingling An, PhD. All experimental procedures with the PGTC were completed in the Mars-Lunar Greenhouse Laboratory at the University of Arizona Controlled Environment Agriculture Center.

## Author contributions

The experimental setup and scope of the project were the result of interactions between MRB, GAG, and HM. QDs were optimized and manufactured by KR. The QD films were optimized and manufactured by AJ. The experiments were planned by MRB, GAG, CHP, and HM. The spectroscopic characterization of the QD films was performed by MRB, AJ, and DH. Modeling was conducted by MRB. The design, operations, and plant trials in the PGTC to obtain the spectroradiometer measurements were completed by CHP and GAG. Data was analyzed by CHP, GAG, MRB, and DH. The manuscript was written by CHP, GAG, and MRB, in consultation with all the authors.

## Competing interests

The authors declare the following competing interests: Authors CHP, DH, AJ, KR, HM, and MRB are employed by UbiQD, Inc. and also have equity in the company. Authors GAG and CHP were not employed by UbiQD, Inc. and had no equity in the company while conducting these experiments.

## Notes

### Summary of Updates

Modeling data has been included in the results section. Supplemental files updated.

