## Supplemental Information for "Optimizing spectral quality with quantum dots to enhance crop yield in controlled environments"

AUTHOR INFORMATION

**Corresponding Authors**

#### Light Uniformity

A metal-halide (MH) lamp (400-W 10K Finisher Lamp, Solis-Tek) was selected as the light source because its radiation emission spectrum is most similar to the solar spectrum of commercially available high-intensity discharge (HID) lamps. The lamp construction consisted of low-iron glass, allowing greater amounts of blue (B) and ultraviolet light (UV) to be discharged from the lamp than typical MH lamps. Four MH lamps of this model were arranged on the upper platform (91 cm above the films) to maximize the uniformity (U) of light intensity incident to all three films. Prior to experimentation, PAR mapping was performed by measuring PPFD (µmol $m^{-2}$ $s^{-1}$) with a full-spectrum quantum sensor (Apogee SQ-500-SS, error < 0.8%) at a height of 12 cm above the test films in five locations as shown in Figure S1. The test films included the Red quantum dot (R-QD), Orange QD (O-QD), and Control (C) films.

##### Uniformity Above Test Films

To ensure that each film received similar quantities of light, PPFD measurements were recorded at quantum sensor height (12 cm) above film level. These measurements and their locations are depicted in the following top-down view relative to the plant locations, denoted by squares, in the chambers below. Open squares symbolize data plant locations, whereas the shaded squares represent guard plants that shielded data plants from edge effects.

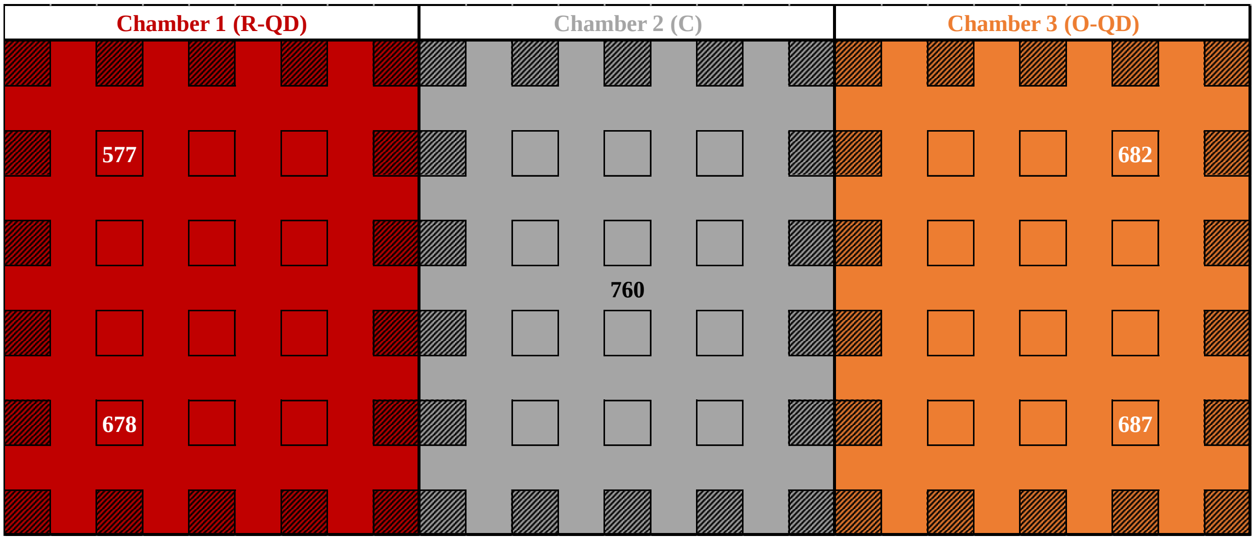

**Supplemental Figure 1.** Top-down view of PAR map of PPFD (µmol $m^{-2}$ $s^{-1}$) before Experiment 1 at five locations above the test films in the PGTC. The R-QD film received the lowest PPFD, while the C film received the greatest.

To determine the light uniformity (U) above the films, the average PPFD ($PPFD_{ave}$=677 µmol $m^{-2}$ $s^{-1}$) was divided by the maximum PPFD ($PPFD_{max}$=760 µmol $m^{-2}$ $s^{-1}$) per the equation:

$$U = \frac{PPFD_{ave}}{PPFD_{max}} = \frac{677\ \mu\text{mol m}^{-2}\text{s}^{-1}}{760\ \mu\text{mol m}^{-2}\text{s}^{-1}} = 0.891$$

#### Uniformity Below Test Films

PPFD measurements were recorded at quantum sensor height (12 cm) above the planting trays and are depicted superimposed over the experimental plant locations, denoted by squares. U values were calculated for each chamber to ensure that plants in each chamber would receive similar levels of light under the films.

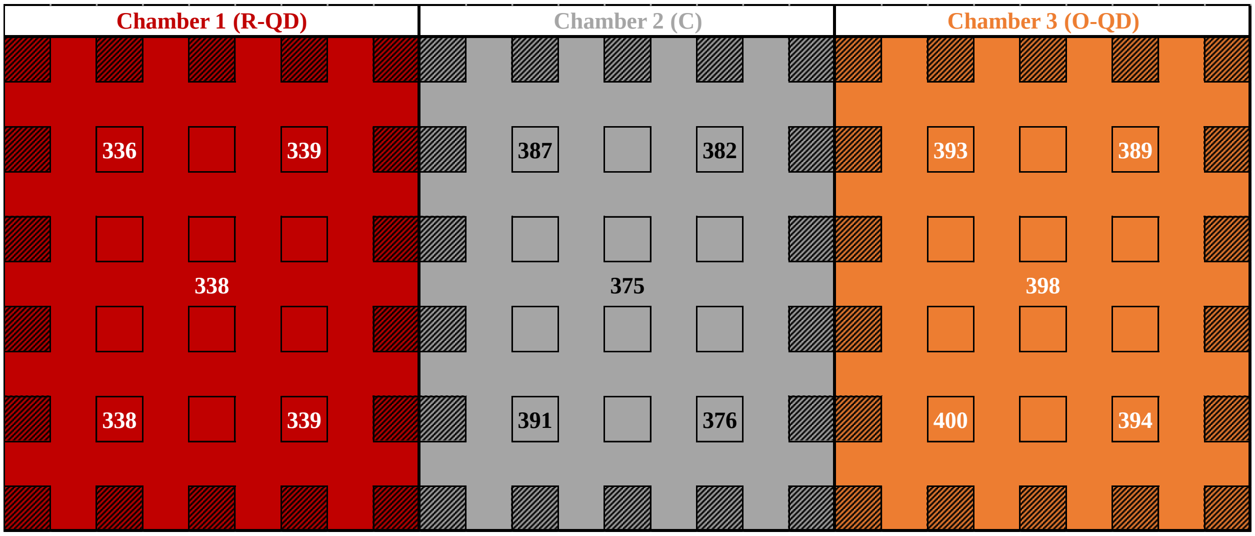

**Supplemental Figure 2.** Top-down view of PAR map of PPFD (μmol m$^{-2}$ s$^{-1}$) before Experiment 1 at five locations below each test film in the PGTC.

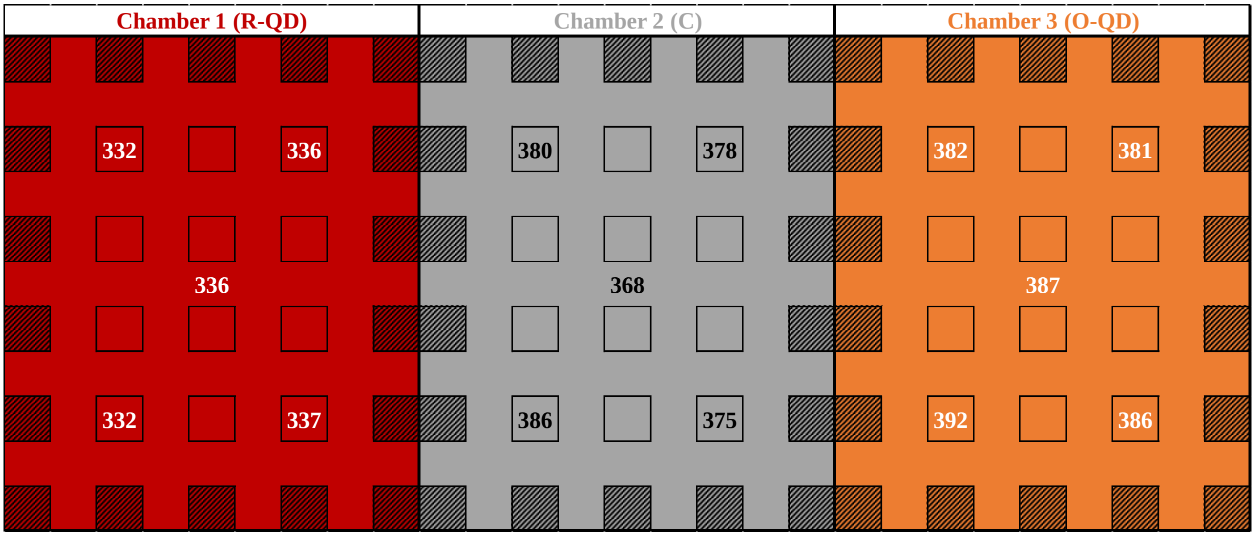

**Supplemental Figure 3.** Top-down view of PAR map of PPFD (μmol m$^{-2}$ s$^{-1}$) before Experiment 2 at five locations below each test film in the PGTC.

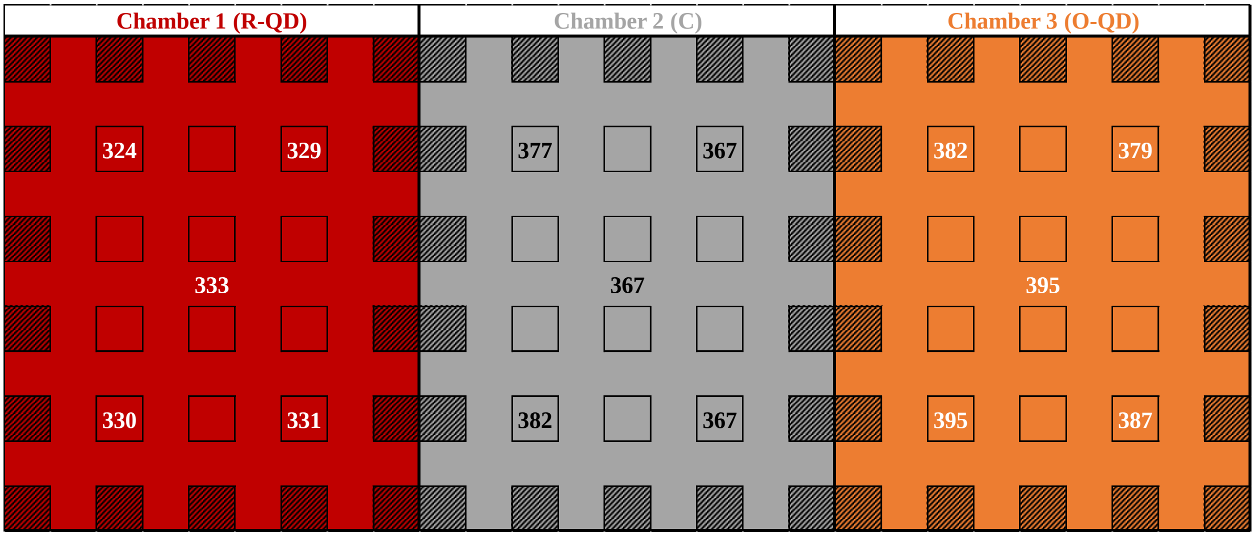

**Supplemental Figure 4.** Top-down view of PAR map of PPFD (μmol $m^{-2}$ $s^{-1}$) before Experiment 3 at five locations below each test film in the PGTC.

**Supplemental Table 1.** PPFD and Uniformity (U) values above and below the test films before Experiment 1.

| Location | Above Films | Below R-QD | Below C | Below O-QD |
|---|---|---|---|---|
| $PPFD_{ave}$ (μmol $m^{-2}$ $s^{-1}$) | 667 | 338 | 382 | 394 |
| $PPFD_{max}$ (μmol $m^{-2}$ $s^{-1}$) | 760 | 339 | 391 | 400 |
| U (%) | 89 | 100 | 98 | 99 |

Prior to experimentation, the U values above and the percentage differences in $PPFD_{ave}$ were compared as shown in Table S1. U values were 98-100%, whereas the PPFD under the C and O-QD films were approximately equal with a difference of 3.1% before Experiment 1. These values were recorded prior to planting, so the absolute values are greater while the differences are lower than those shown in the Light Use Efficiency section.

#### Daily Light Integral Calculations

To calculate the daily light integral (DLI, mol $m^{-2}$ $d^{-1}$), 15-minute $PPFD_{ave}$ measurements from the quantum sensors (Apogee Instruments SQ-500-SS) located at plant canopy height in each treatment zone were recorded for experiments 1-3. Throughout each experiment, the PPFD values gradually decreased due to plant canopy closure reducing the albedo within each chamber. Between each experiment, the $PPFD_{ave}$ values decreased 5-9% due to aging of the metal-halide lamps, but the ratios of the PPFD values under the QD films to the Control Film remained approximately constant. The average DLI values ($DLI_{ave}$), shown in Table S2 for each treatment group per experiment, were calculated by taking the sum of all $PPFD_{ave}$ values over each 14-h photoperiod with the following equation:

$$DLI_{ave} = \sum PPFD_{ave}$$

**Supplemental Table 2.** DLI measurements at plant canopy height in each plant experiment and $DLI_{ave}$ values for each treatment across all three experiments

| | **Experiment 1** | | **Experiment 2** | | **Experiment 3** | | **Experiments 1-3** | |
|---|---|---|---|---|---|---|---|---|
| **Film** | **$DLI_{ave}$ (mol $m^{-2}$ $d^{-1}$)** | **Diff. v. C (%)** | **$DLI_{ave}$ (mol $m^{-2}$ $d^{-1}$)** | **Diff. v. C (%)** | **$DLI_{ave}$ (mol $m^{-2}$ $d^{-1}$)** | **Diff. v. C (%)** | **$DLI_{ave}$ (mol $m^{-2}$ $d^{-1}$)** | **Diff. v. C (%)** |
| Control | 17.9 | - | 16.6 (Δ=-1.3) | - | 16.6 (Δ=0.0) | - | 17.0 | **-** |
| O-QD | 19.0 | +6.0 | 18.0 (Δ=-1.0) | +8.3 | 17.6 (Δ=-0.4) | +5.9 | 18.2 | +6.7 |
| R-QD | 16.0 | -11.0 | 15.2 (Δ=-0.8) | -8.4 | 14.9 (Δ=-0.3) | -10.3 | 15.4 | -9.9 |

The above DLI values were used to calculate the light use efficiency (LUE, g $mol^{-1}$), as seen in the subsequent section, but not the percentage difference in incident light provided to the plants (See Light Uniformity section above).

In Figures S5-S7 below, the measured DLI decreased after 14 DAS in each experiment due to more photons being absorbed by the plant canopy and fewer photons being reflected off the white base of the planting area as the plant canopy closed in each chamber. The following graphs are based on instantaneous quantum sensors readings in the presence of plants and so are affected by the reduced albedo in the chambers.

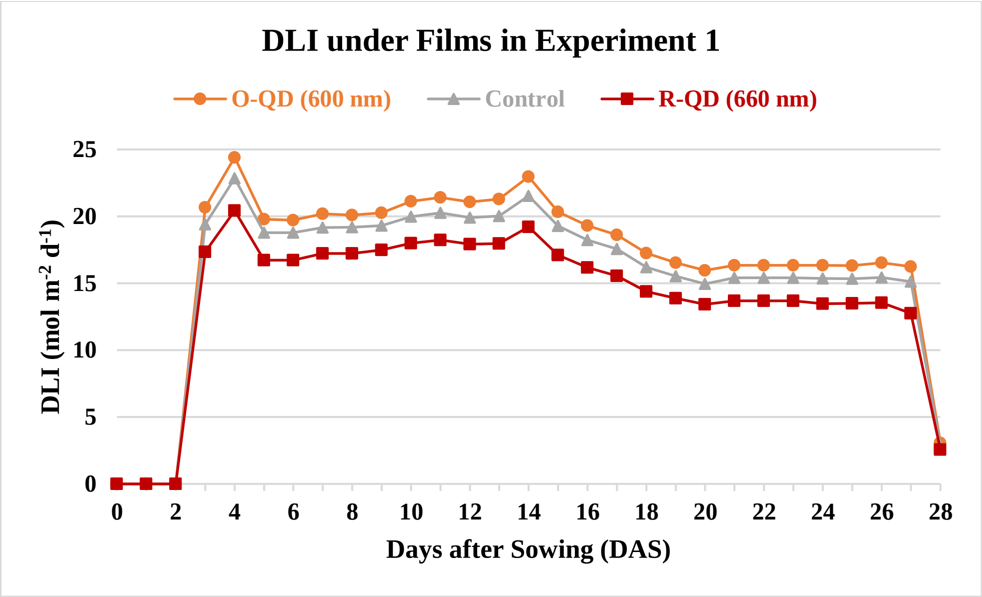

**Supplemental Figure 5.** Daily Light Integral (DLI) values under each film in Experiment 1. Lighting began upon observation of germination 3 DAS. Spikes at 4 DAS and 14 DAS were due to PGTC system inspection requiring additional lighting time.

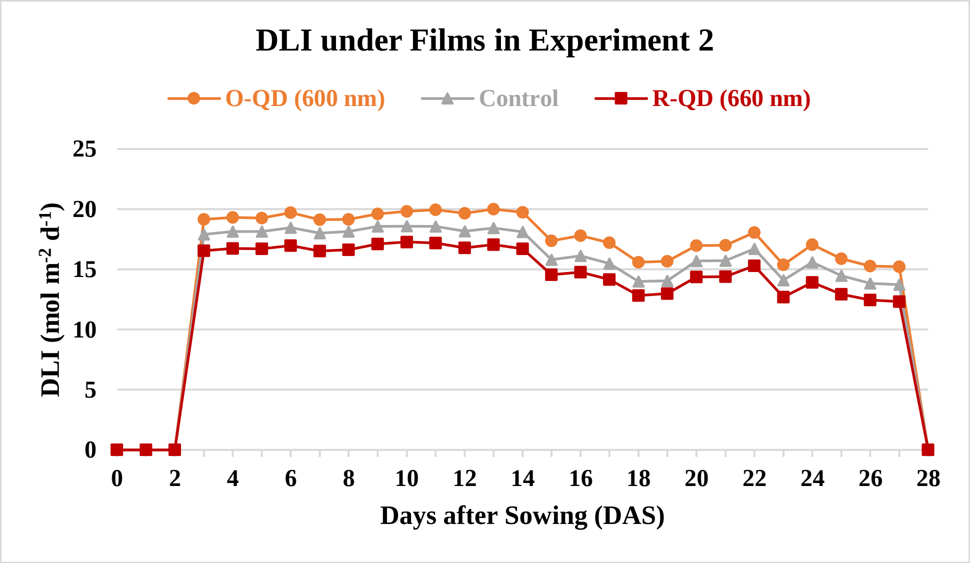

**Supplemental Figure 6.** Daily Light Integral (DLI) values under each film in Experiment 2. Lighting began upon observation of germination 3 DAS. Relatively low measurements after 17 DAS were due to quantum sensor coverage by the leaf canopy.

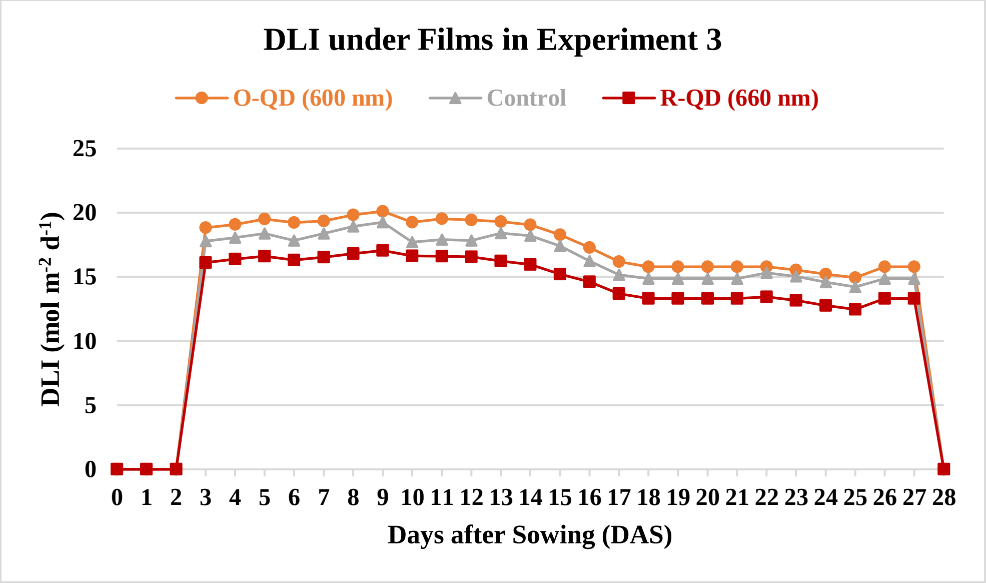

**Supplemental Figure 7.** Daily Light Integral (DLI) values under each film in Experiment 3. Lighting began upon observation of germination 3 DAS. Relatively low measurements after 24 DAS were due to quantum sensor coverage by the leaf canopy.

#### Light Use Efficiency

The LUE represents the biomass production per DLI given to the plant, which can be calculated from fresh mass (FM) or dry mass (DM) measurements. The LUE values were calculated by taking the ratio of the average daily mass production per area to the average DLI ($DLI_{ave}$). The LUE was calculated for the $FM_{ave}$ (including water weight) and $DM_{ave}$, which allowed for comparison of the production efficiencies between the three light treatments.

After calculating the DLIs, the average daily mass productions per area were calculated for each of the three film treatments. The average total FM was 44.8 g, 45.6 g, and 40.9 g under the R-QD, O-QD, and C films, respectively. The average total DM was 1.55 g, 1.60 g, and 1.42 g under the R-QD, O-QD, and C films, respectively. The mass production per area (g $m^{-2}$) was calculated for each zone by dividing the average mass production by the number of harvested plants (n=12) and multiplying by the planting density (55 plants $m^{-2}$). The production occurred over a period of 28 days, so the mass production per area was divided by the experiment duration to yield the average daily FM ($FM_{ave}$) production (g $m^{-2}$ $d^{-1}$), displayed in Table S3.

From these calculations, the R-QD Film LUE was ~17% greater for FM and ~18% greater for DM than the C Film, while the O-QD Film LUE was ~4% greater for FM and 6% greater for DM than the C Film. The greater LUE values under both QD films clearly indicate that photosynthetic efficiency was increased by improving the spectral quality.

**Supplemental Table 3.** Average daily production, average DLI, and calculated LUE across all experiments for three different film treatments. Percentage differences in LUE are under QD Film v. Control (C) Film.

| | **R-QD Film** | **O-QD Film** | **C Film** |
|---|---|---|---|
| **Daily $FM_{ave}$ Production** (g $m^{-2}$ $d^{-1}$) | 7.33 | 7.46 | 6.70 |
| **Daily $DM_{ave}$ Production** (g $m^{-2}$ $d^{-1}$) | 0.254 | 0.262 | 0.232 |
| **$DLI_{ave}$** (mol $m^{-2}$ $d^{-1}$) | 15.4 | 18.2 | 17.0 |
| **$FM_{ave}$ LUE** (g $mol^{-1}$) | 0.476 **(+17.2% v. C)** | 0.410 **(+3.90% v. C)** | 0.394 |
| **$DM_{ave}$ LUE** (g $mol^{-1}$) | 0.0165 **(+17.6% v. C)** | 0.0144 **(+5.55% v. C)** | 0.0136 |

#### Total Leaf Area Analysis Program

As a leaf area meter was not available for use, a software program was written in C++ and used Open Graphics Library (OpenGL) and OpenGL Shader Language (GLSL) for total leaf area (TLA) measurement. The program was designed to render a flat texture containing the image of the leaves and apply a fragment shader and a color-based filter to remove non-leaf pixels from the image. The program also employed keystone correction prior to image analysis to correct area distortions introduced by slight misalignment between the image capture stage and the image focal plane. After the leaves had been selected from the background, the program performed a calibrated area measurement based on a known reference area included in each photograph (Figure S8).

To validate the accuracy of the program measurements, validation photographs were taken using an object of a known area, and the program reliably calculated TLA of these images with <1% error. A reference area was included in each leaf photograph; a 36-$cm^2$ paper was included to further increase the reference area and thereby reduce measurement error. For most images, the camera position was fixed such that only a 1-pixel error was possible in the reference area, so the total error was not likely to exceed the above values plus 1% (the estimated combined error from other sources), resulting in a total error of <2% for most images to <5% for the first set in Experiment 1. The largest potential source of error was the estimation of the pixels in the reference area, so estimation was performed by a single operator, to minimize variation in area estimation. The operator also performed manual corrections to the images, as needed, to improve the filter's accuracy in leaf selection.

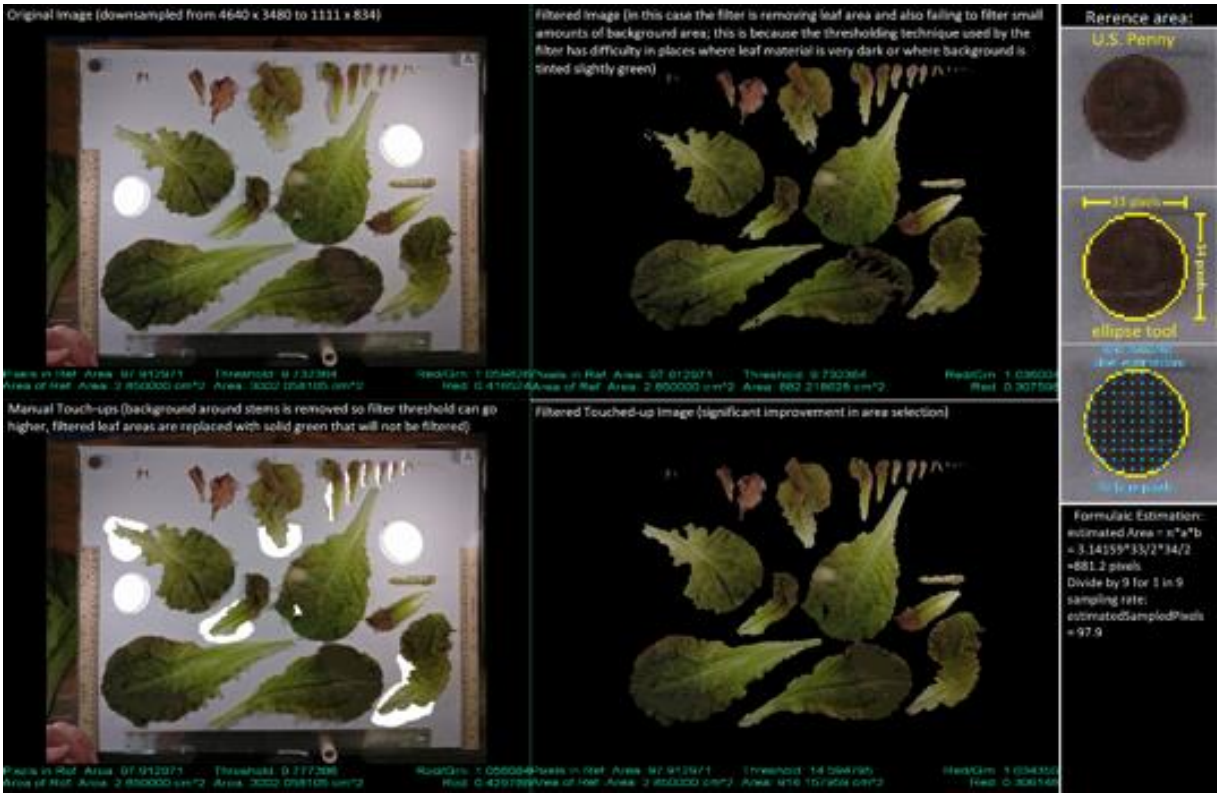

**Supplemental Figure 8.** Example of software program performing leaf area analysis based on reference area of a U.S. penny. The reference area was subsequently increased to a 36-$cm^2$ area to reduce error in total leaf area (TLA) calculations to <1%.

#### Dry Mass Percentage Calculation

The edible dry mass (DM) percentage (DM%) was calculated for the plant material resulting from each experiment. Individually bagged FM samples from each experiment were dried at 40 °C for >72 h to remove all water content and to achieve consistently uniform DM% within each experiment to ensure comparability within and between experiments.

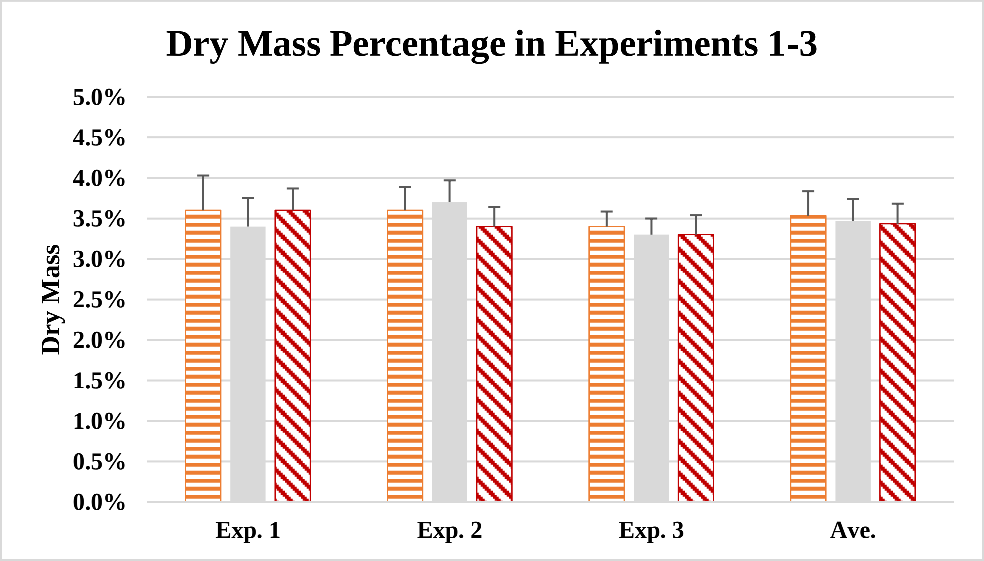

**Supplemental Figure 9**. Dry mass (DM) percentages of fresh mass (FM) in each experiment. Average DM percentage across all experiments was 3.5% for each film.

### Environmental Parameters

#### pH & Electrical Conductivity (EC) of Shared Hydroponic Nutrient Solution

The automated pH controller performed 'hunting' for the set point between 6.1-6.2, and this effect was more pronounced prior to the transition from tap water to half-strength modified Hoagland's solution upon germination at 3 DAS. Transition from half- to full-strength solution occurred at 7 DAS and was maintained manually between 1.8-1.9 $mS^{-1}$ with daily additions of concentrated nutrient solution and/or tap water.

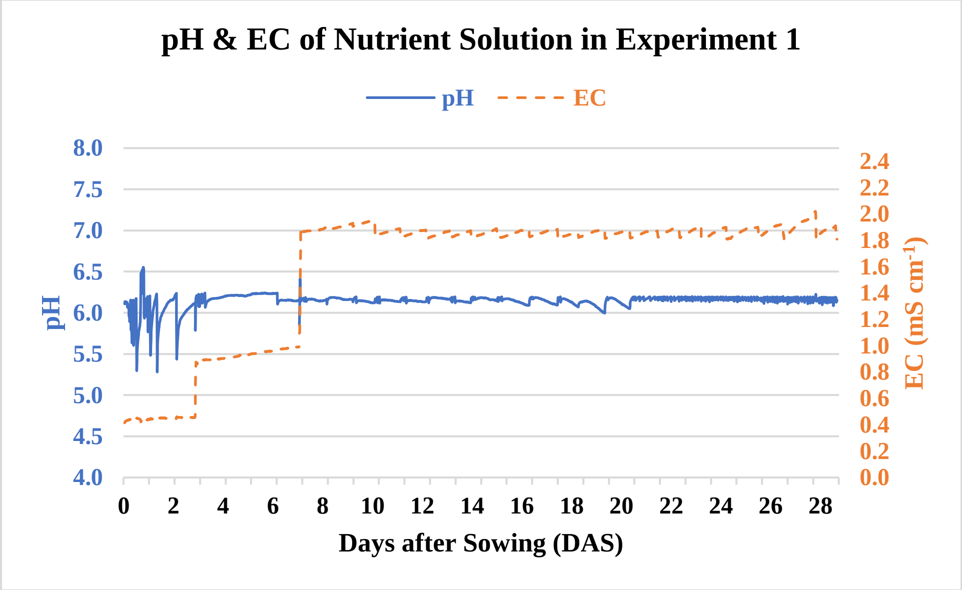

**Supplemental Figure 10.** pH and electrical conductivity (EC) values of the shared hydroponic nutrient solution in Experiment 1.

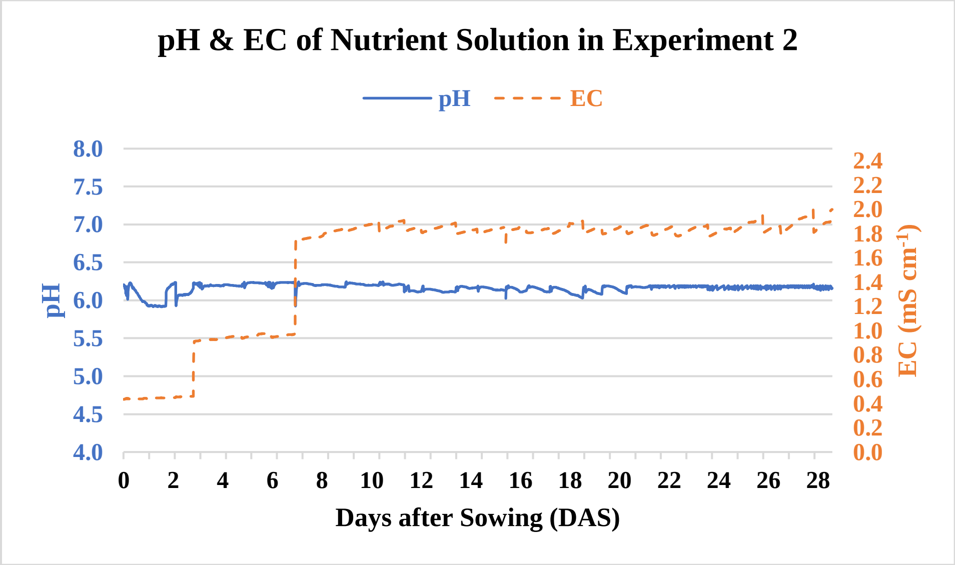

**Supplemental Figure 11.** pH and electrical conductivity (EC) values of the shared hydroponic nutrient solution in Experiment 2.

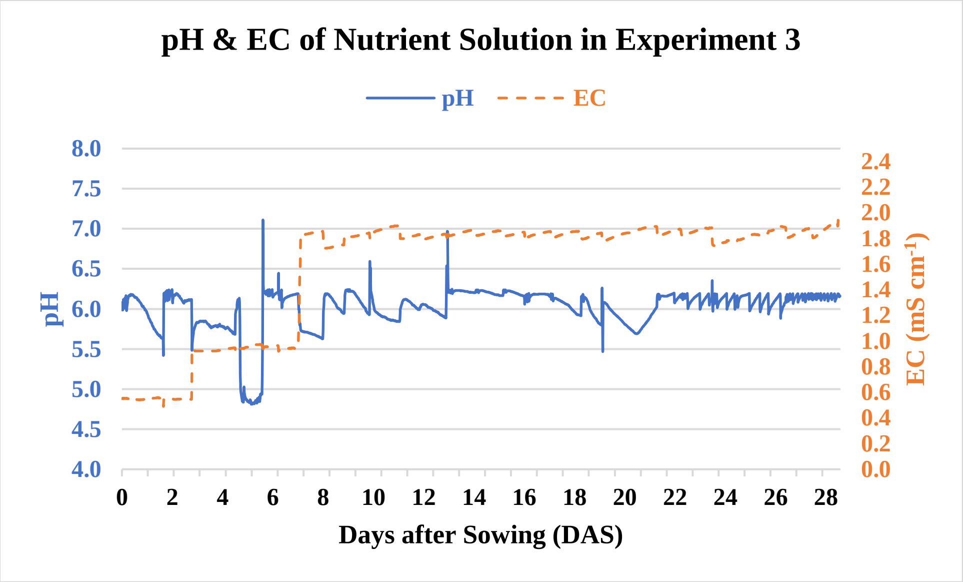

**Supplemental Figure 12.** pH and electrical conductivity (EC) values of the shared hydroponic nutrient solution in Experiment 3. The peaks and troughs in the pH readings were due to the automated peristaltic pump overdosing concentrated nitric acid and resulting manual addition of tap water to return solution to set point.

#### Temperature of Plant Chambers and Root Zone

15-minute average temperature readings were recorded throughout each experiment. These temperatures included that of the air at plant canopy height in each of the three test chambers as well as the root zone temperature of the shared hydroponic nutrient solution. Temperatures increased after lighting began at 3 DAS, when a photoperiod of 14 h $d^{-1}$ was applied.

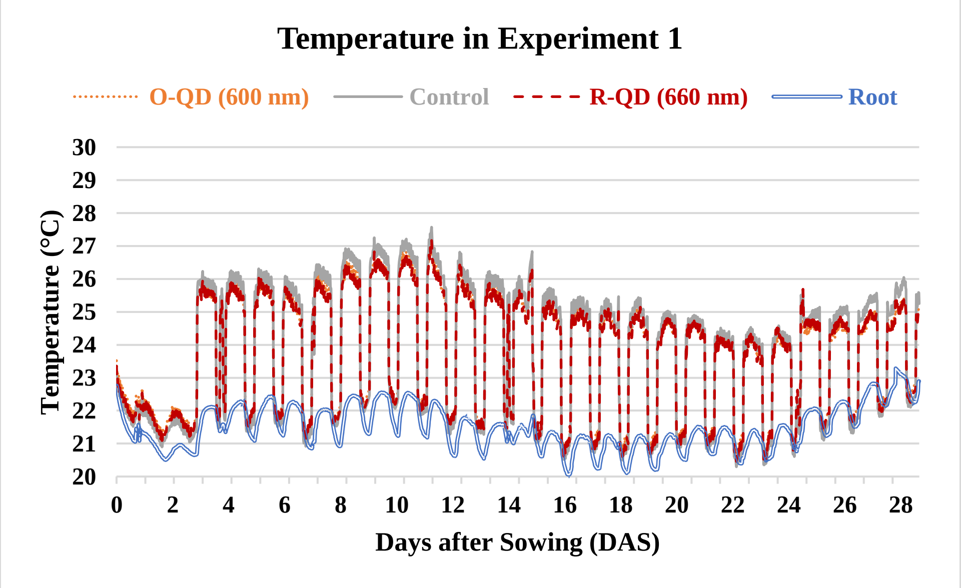

**Supplemental Figure 13.** Temperatures at plant canopy height within each test chamber as well as temperature of the root zone in the shared hydroponic nutrient solution in Experiment 1. Average air temperature differences between treatments remained within ±1 °C for the duration of the experiment.

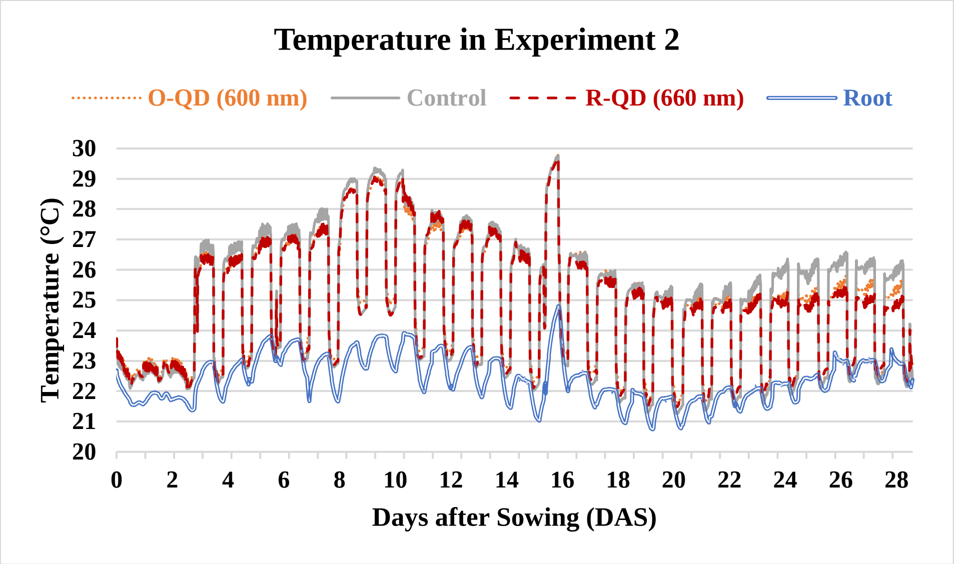

**Supplemental Figure 14.** Temperatures at plant canopy height within each test chamber as well as temperature of the root zone in the shared hydroponic nutrient solution in Experiment 2. The spike at 17 DAS occurred on a particularly hot day in Tucson, AZ, when the laboratory air conditioning could not adequately compensate. Average air temperature differences between treatments remained within ±1 °C for the duration of the experiment but widened between the Control Film and the two QD films after 23 DAS.

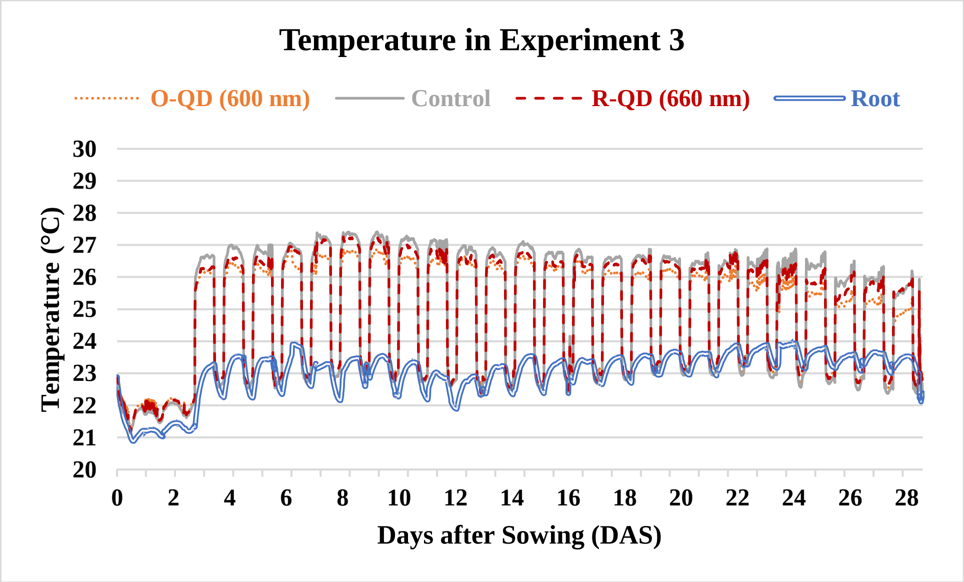

**Supplemental Figure 15.** Temperatures at plant canopy height within each test chamber as well as temperature of the root zone in the shared hydroponic nutrient solution in Experiment 3. Average air temperature differences between treatments remained within ±1 °C for the duration of the experiment.

**Carbon Dioxide ($CO_2$) Concentration**

Average $CO_2$ concentration remained between 350 ppm and 400 ppm for all three experiments. Spikes in the sensor readings occurred when human operators were present performing daily operations and routine maintenance on the PGTC in the laboratory.

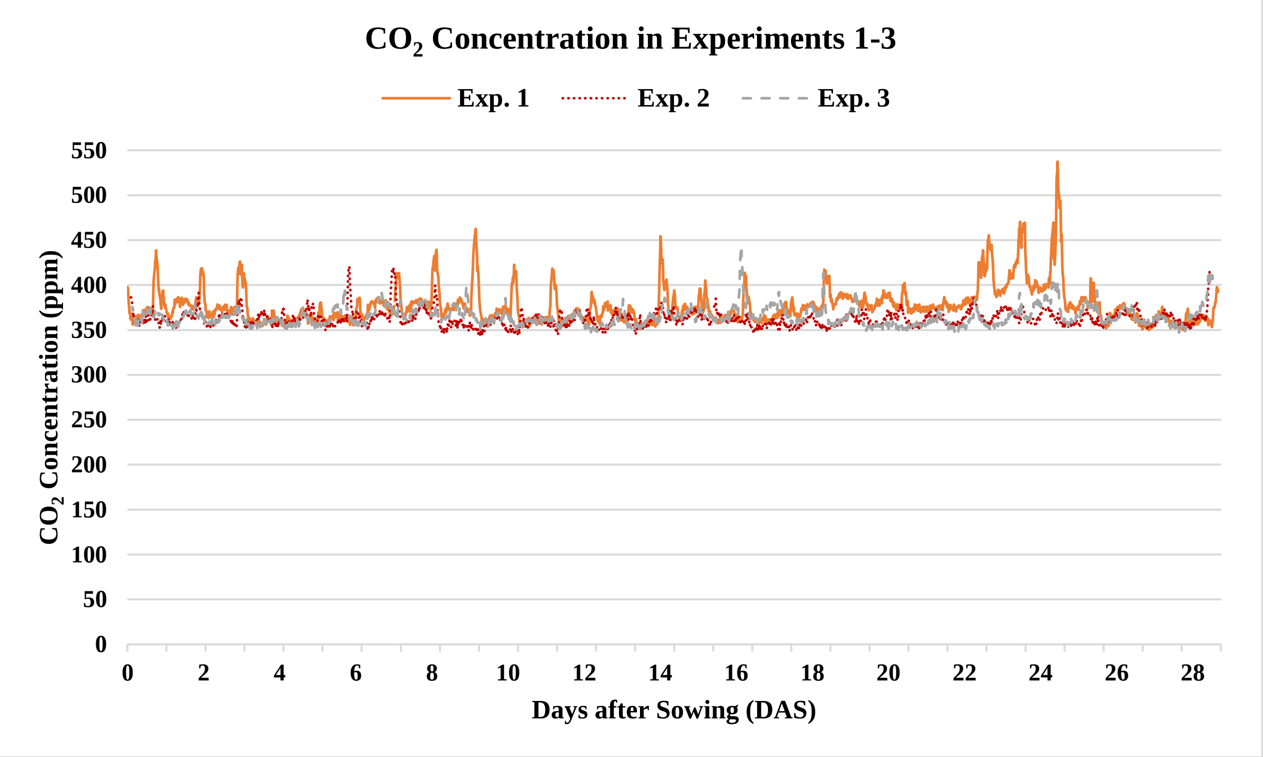

**Supplemental Figure 16.** Carbon dioxide ($CO_2$) concentration in the room during Experiments 1-3 remained 350-400 ppm, except for spikes in the sensor readings when human operators were present performing daily operations and routine maintenance near the PGTC in the laboratory.

#### Plant Layout

Thirty plants were located under each of the three films in each experiment with a quantum sensor placed at a central location at plant canopy height in each chamber. The 18 guard plants as shown in Figure S14 were planted to reduce edge effects observed in the 12 data plants, which were the only plants measured.

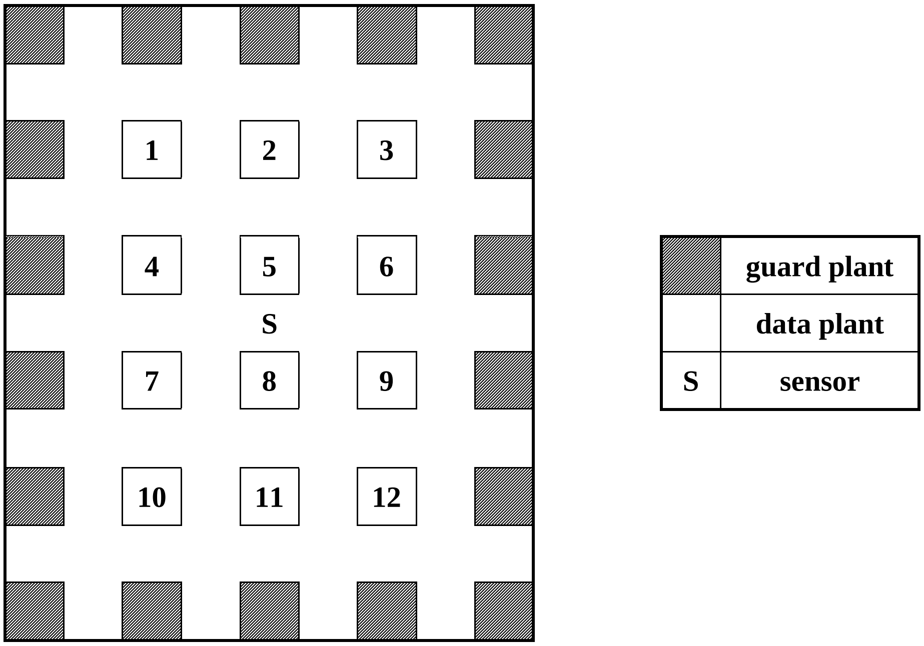

**Supplemental Figure 17**. Plant arrangement under each film. In each experiment, the total number of data plants per treatment group was n=12; thus, after three experiments, N=36 total replicates per treatment group were achieved. Plants were spaced 12 cm center-to-center in each row and column at a density of 55 plants $m^{-2}$. Throughout each experiment, a quantum sensor was placed in the center of the grow area under each film.

**Supplemental Table 4**. Lettuce nutrient formula. Approximately half-strength modified Hoagland's solution.

| | | | | | | | | |
|---|---|---|---|---|---|---|---|---|
| Recommended Lettuce Nutrient Solution (CEAC, 2/22/2008) | | | | | | | | |
| Assumptions: | | | | | | | | |
| Two tanks - 25 gal. (95 L) each | | | | | | | | |
| Fertilizer injector ratio = 100:1 | | | | | | | | |
| | | Elements (ppm) | | | | | | |
| Tank A | | N | P | K | Ca | Mg | S | Cl |
| Ca(NO3)2 | | 134 | | | 165 | | | |
| CaCl2.6H2O | | | | | 35 | | | 60 |
| Tank B | | | | | | | | |
| KNO3 | | 46 | | 137 | | | | |
| KH2PO4 | | | 50 | 61 | | | | |
| MgSO4.7H2O | | | | | | 40 | 52 | |
| Total ppm | | 180 | 50 | 198 | 200 | 40 | 52 | 60 |
| Macroelements | | | | | | | | |
| Tank A | | | | | | | | |
| Ca(NO3)2 | | | 8.2 | kg | | | | |
| CaCl2.6H2O | | | 1.8 | kg | | | | |
| | | Total | 10.0 | kg | | | | |
| Tank B | | | | | | | | |
| KNO3 | | | 3.4 | kg | | | | |
| KH2PO4 | | | 2.0 | kg | | | | |
| MgSO4.7H2O | | | 3.8 | kg | | | | |
| | | Total | 9.2 | kg | | | | |
| Microelements | | | | | | | | |
| Tank A | | | | | | | | |
| Iron chelate EDTA | | (13.2%)Fe | 175.0 | g | 2.4 | ppm | | |
| Tank B | | | | | | | | |
| Manganese sulfate | | **(32%)Mn** | 16.00 | g | 0.55 | ppm | | |
| Zinc sulfate | | (36%)Zn | 9.00 | g | 0.34 | ppm | | |
| Copper sulfate | | (25%)Cu | 1.90 | g | 0.05 | ppm | | |
| Solubor | | (20.5%)B | 16.00 | g | 0.35 | ppm | | |
| Sodium molybdate | | (40%)Mo | 1.20 | g | 0.05 | ppm | | |
| | | Total | 44.10 | g | 1.34 | ppm | | |

#### Spectra

Absorption measurements on the O-QD and R-QD films were made using a Agilent Technology Cary 8454 UV-Vis absorption spectrometer and photoluminescence spectra were measured on a Horiba Fluoromax-4 emission spectrometer. Figures S15 and S16 illustrate the Stokes shift between peak emission and absorption.

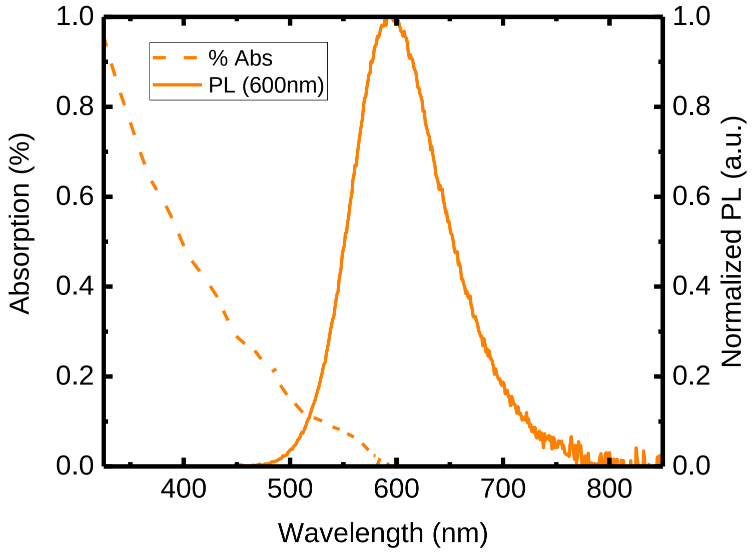

**Supplemental Figure 18**. Percent absorption (dashed line) and normalized PL emission (solid line) for O-QD film are plotted illustrating the Stokes shift between the peak emission and absorption by CIS/ZnS QDs.

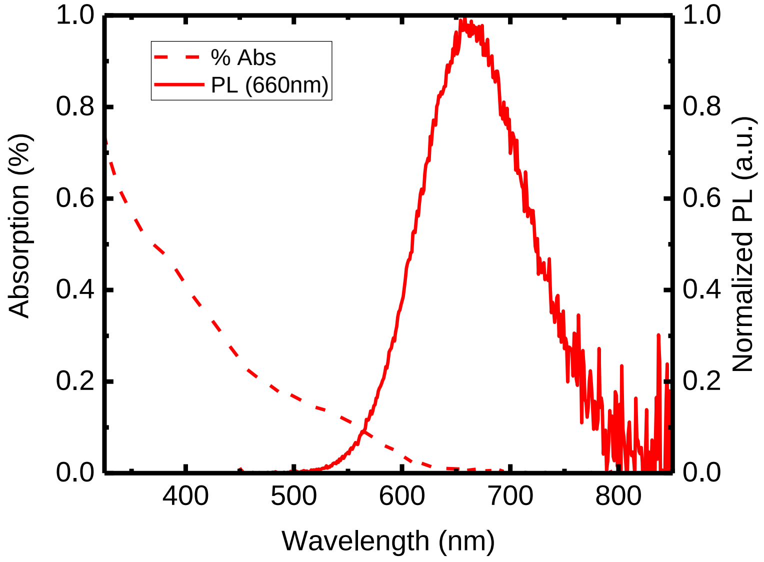

**Supplemental Figure 19**. Percent absorption (dashed line) and normalized PL emission (solid line) for R-QD film are plotted.

PPFD spectra were measured underneath the O-QD film, control film, and R-QD film with an SRI-PL-6000 Spectrophotometer (Optimum Optoelectronics Corp.) and comparisons of spectrum between QD film and control film are shown below.

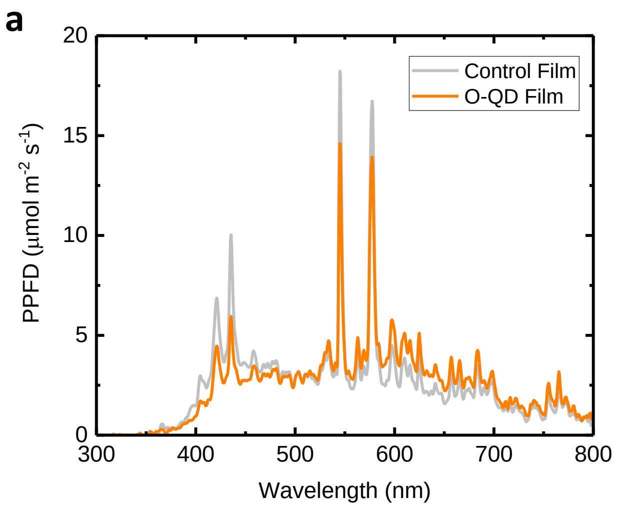

**Supplemental Figure 20.** Measured spectra for each of the three light treatments. (a) Measured spectra beneath the O-QD film and under the control. (b) Measured spectra beneath the R-QD film and control film.

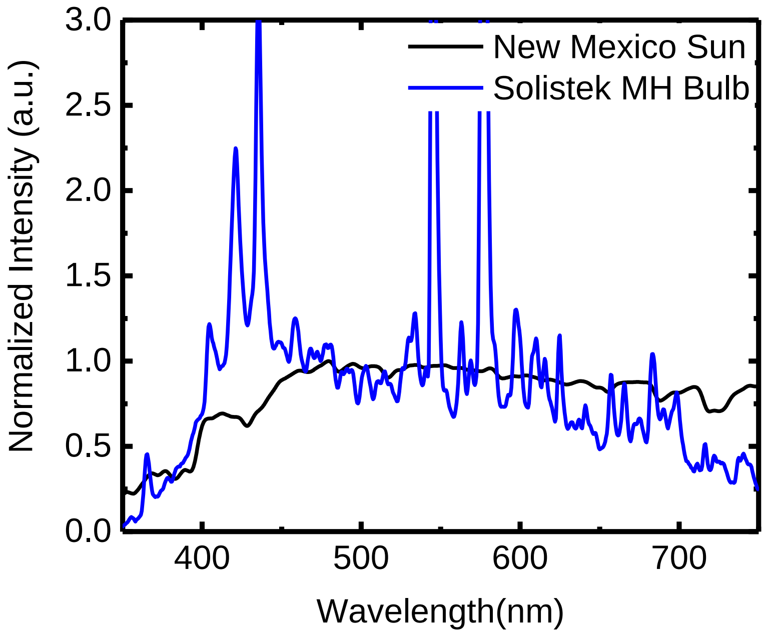

**Supplemental Figure 21.** Comparison of the metal-halide (MH) lamp model spectra with solar spectra. Spectra were measured by an SRI-PL-6000 portable spectrometer and normalized at 475nm. Solar spectrum was measured at noon on October 10, 2018 in Los Alamos, New Mexico.

### Modeling

First, the solar irradiance values for reference air mass (AM) 1.5 and 1.0 spectra from NREL1 were converted from W m-2 nm-1, to solar photon flux density (Figure S22). To convert the spectra, first, the number of photons (NP) per second per area is calculated from the irradiance (I) using the following equation: $N_p = I/(\frac{hc}{\lambda})$ , where $\frac{hc}{\lambda}$ is the energy of a photon of a given wavelength ($\lambda$). Then the spectra are divided by Avogadro's number ($N_p = 6.02x10^{-7}\ \mu mol^{-1}$) to get the photon flux in units of µmol $m^{-2}$ $s^{-1}$.

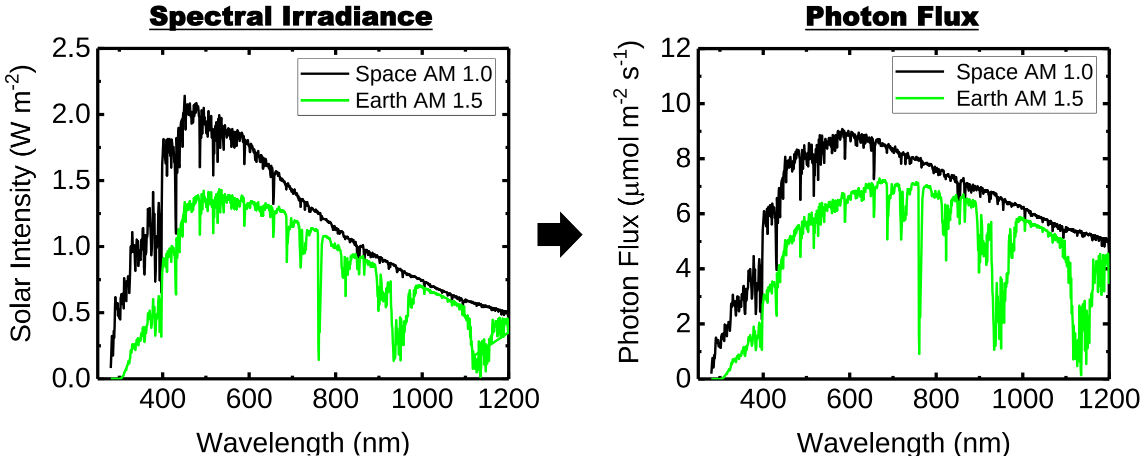

**Supplement Figure 22.** Conversion of solar irradiance data of reference AM 1.5 spectra (terrestrial) and AM 1.0 spectra (space) to solar photon flux density.

Once the two spectra are converted to photon flux, then the % absorption spectra for both the O-QD and R-QD films were convoluted with both solar spectra (Supplemental Figure 23 b,c). Supplemental Table 5 shows the calculated results for each wavelength range under both QD films for both AM 1.0 and AM 1.5 spectra.

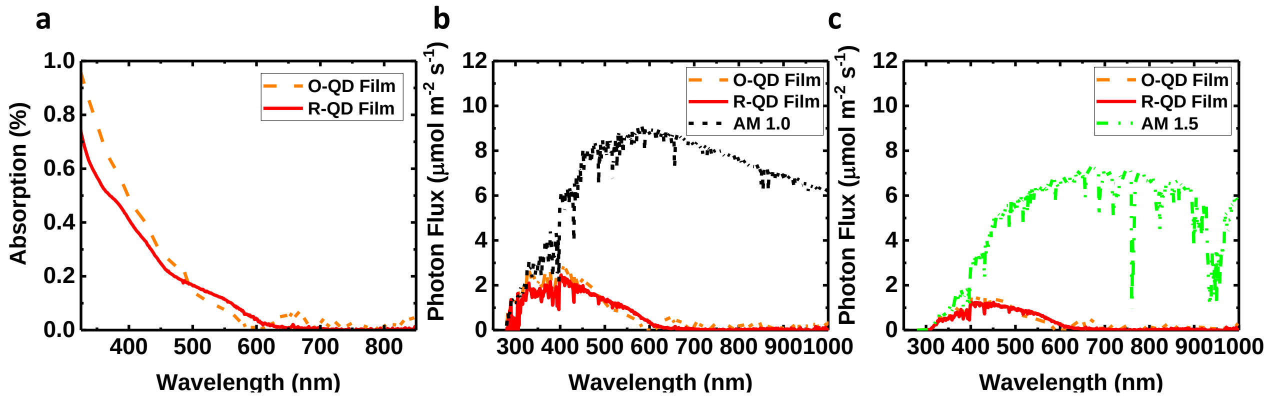

**Supplemental Figure 23.** (a) Absorption spectrum of the O-QD (600 nm) and R-QD (660 nm) Films in terms of percent absorption per wavelength. Convolution of the absorption spectra of the O-QD and R-QD films with the AM 1.0 spectra (b) and the AM1.5 spectra (c).

**Supplemental Table 5.** QD Film absorption calculations for AM 1.0 and AM 1.5 solar spectrum.

| **AM 1.0** | | | | | | | | | | |
|---|---|---|---|---|---|---|---|---|---|---|
| **Film** | **Total PPFD (µmol $m^{-2}$ $s^{-1}$) 400-700 nm** | **UV PFD (µmol $m^{-2}$ $s^{-1}$) <400 nm** | **UV Abs (µmol $m^{-2}$ $s^{-1}$)** | **UV Abs (%)** | **B PFD (µmol $m^{-2}$ $s^{-1}$) 400-500 nm** | **B Abs (µmol $m^{-2}$ $s^{-1}$)** | **B Abs (%)** | **G PFD (µmol $m^{-2}$ $s^{-1}$) 500-600 nm** | **G Abs (µmol $m^{-2}$ $s^{-1}$)** | **G Abs (%)** |
| O-QD | 2413.2 | 301.1 | 290.0 | 96.4% | 704.4 | 432.4 | 61.4% | 848.4 | 76.8 | 9.1% |
| R-QD | 2413.2 | 301.1 | 276.6 | 91.9% | 704.4 | 345.7 | 49.1% | 848.4 | 102.2 | 12.1% |
| **AM 1.5** | | | | | | | | | | |
| O-QD | 1735.3 | 93.4 | 88.7 | 94.9% | 437.3 | 262.3 | 60.0% | 615.0 | 54.8 | 8.9% |
| R-QD | 1735.3 | 93.4 | 82.9 | 88.7% | 437.3 | 209.1 | 47.8% | 615.0 | 73.3 | 11.9% |

[1] Reference Solar Spectral Irradiance: Air Mass 1.5" National Renewable Energy Laboratory. Retrieved from https://rredc.nrel.gov/solar//spectra/am1.5/
